# Capsanthin/Capsorubin Synthase Expression in Tomato Alters Carotenoid Pools, Enhancing Provitamin A and Flavor Volatiles

**DOI:** 10.1101/2024.09.27.615503

**Authors:** Jingwei Fu, Denise Tieman, Bala Rathinasabapathi

**Author notes:** **One sentence summary** Expression of capsanthin/capsorubin synthase in tomato enhanced total carotenoids, β-carotene, ketocarotenoids and flavor volatiles in ripening fruit with no impact on plant growth and development.

## Abstract

We conceptualized a tomato biofortification strategy via engineering simultaneous accumulation of *β*-carotene, a provitamin A and pepper-specialized ketocarotenoids, capsanthin, and capsorubin. Capsanthin/capsorubin synthase (CCS) in pepper, an enzyme phylogenetically related to lycopene β-cyclases (LCYB) known for *β*-carotene synthesis, was investigated for its *in vivo* role in ripening fruit. In pepper, silencing of *CCS* via virus-induced gene silencing reduced the flux from lycopene to *β*-carotene (the *β*-branch) with significant changes in carotenoid compositions. In a carotenogenic bacterial expression system, pepper CCS was more effective than tomato LCYB1 or LCYB2 in lycopene cyclization to *β*-carotene and CCS synthesized capsanthin, but the LCYBs did not. Therefore, we overexpressed pepper *CCS* in different tomato varieties, including ‘Micro-Tom’ WT, ‘Micro-Tom’ mutant, *pyp1-1(H7L)* (defective in xanthophyll esterification), and five inbreds and characterized their carotenoid profiles. In *CCS*-transformed tomato fruit (WT and selected varieties), besides the biosynthesis of capsanthin and capsorubin, total carotenoids, *β*-carotene, and xanthophyll esters remarkably increased compared to the controls, while such increments were weaker in the *pyp1-1(H7L)*. CCS expression had a positive influence on the flux toward the *β*-branch and the storage pool of xanthophyll esters consistent with its dual functions in lycopene cyclization and ketocarotenoid biosynthesis. The data further supported that xanthophyll esters facilitated carotenoid accumulation. While CCS-expression had no significant impact on growth or yield, fruit of *CCS*-transformed tomato had greater levels of *β*-carotene-derived flavor volatiles than the controls. Consumption of 37–131-gram of *CCS*-derived hybrid fruit meets the provitamin A recommended dietary allowance, indicating greatly improved nutritional value.

## Introduction

Pepper fruit, a nutrient-dense vegetable for human consumption, contains health-promoting metabolites, including vitamins, alkaloids, and specialized metabolites such as capsanthin, and capsorubin (Wahyuni *et al*., 2013). Capsanthin and capsorubin are ketocarotenoids, accumulating in the ripe fruit of *Capsicum* and a few other taxa (Trajković *et al*., 2021). Capsanthin/capsorubin synthase (CCS, EC no.: 5.3.99.8) is responsible for the biosynthesis of capsanthin and capsorubin using antheraxanthin and violaxanthin as substrates, respectively (Bouvier *et al*., 1994; Mialoundama *et al*., 2010). CCS was first identified from pepper (Bouvier *et al*., 1994), and validated through enzymatic assays *in vitro* (Mialoundama *et al*., 2010). In addition to *k*-cyclase activity, CCS was reported to have the catalytic activity of lycopene beta-cyclase (LCYB) also, via *in vitro* assays converting lycopene into *β*-carotene (Cunningham *et al*., 1994; Bouvier *et al*., 1997; Mialoundama *et al*., 2010). In pepper fruit, the majority of capsanthin and capsorubin are esterified by a xanthophyll acyltransferase (XAT) and stored inside the plastoglobule (Berry *et al*., 2019). The xanthophyll esterification to fatty acyl groups improves their accumulation, stability, and storage (Ariizumi *et al*., 2014; Li *et al*., 2023). However, the relationship between the biosynthesis of abundant ketocarotenoids and the process of xanthophyll esterification in pepper fruit is still unclear. The *in vivo* role of CCS in the carotenoid pool of ripe pepper fruit remains poorly understood as well, although CCS was well known to have lycopene cyclase and *k*-cyclase activities and be associated with xanthophyll esterification process. We hypothesized that transferring *CCS* from pepper to other related plant species could allow us to test its role in carotenoid biosynthetic flux and accumulation, and the production of specialized ketocarotenoids, and their esters. To date, stable genetic transformation of plants expressing *CCS* in the endosperm of rice (Ha *et al*., 2019) and in the flower of *Viola cornuta* (Trajković *et al*., 2021) have been reported. However, the alteration of the carotenoid pathway by CCS in a fleshy fruit has remained elusive.

Metabolic engineering is an efficient method to biofortify crops in a targeted direction (Van Der Straeten *et al*., 2020), such as purple tomato engineered to accumulate anthocyanins (Butelli *et al*., 2008). Vitamin A is indispensable for human health since vitamin A deficiency (VAD) may result in the development of disorders like xerophthalmia, blindness, and anemia (Blaner, 2020). Globally, an estimated 250–500 million preschool children suffer from VAD (World Health Organization). Hence, carotenoid biofortification of nutrient-dense plant-based foods is an excellent approach to simultaneously combat food insecurity and VAD (Kraemer *et al*., 2008; Grune *et al*., 2010; Fitzpatrick *et al*., 2012; Blaner, 2020).

Multiple cases of carotenoid pathway engineering have been performed to biofortify crops with *β*-carotene, the major provitamin A (Giuliano, 2017; Zheng *et al*., 2020), including the first example, golden rice, in which *β*-carotene accumulated in rice endosperm (Ye & Beyer, 2000). In addition to provitamin A, the alteration of non-provitamin A carotenoids, like lycopene, lutein, and zeaxanthin, through metabolic engineering is also of interest. The pepper ketocarotenoids, capsanthin and capsorubin, are reported to have higher antioxidant ability and potential health promoting activities tested by cell culture and animal model studies (Nishino *et al*., 2016; Kennedy *et al*., 2021). Besides crops, carotenoid metabolic engineering has been fulfilled in microbes, especially in *E. coli* (Lee & Schmidt-Dannert, 2002; Furubayashi *et al*., 2021; Xu *et al*., 2023).

Hence, we aimed to (1) test the *in vivo* role of CCS in carotenoid pools of pepper fruit, (2) compare enzymatic function of pepper CCS and tomato LCYBs, and (3) test the impact of CCS overexpression and xanthophyll esterification in tomato fruit potentially increasing the levels of total carotenoids, provitamin A carotenoid *β*-carotene, volatile apocarotenoids and ketocarotenoids capsanthin and capsorubin. Taking advantage of multiple functions of CCS, we overexpressed pepper *CCS* in ‘Micro-Tom’ WT and a ‘Micro-Tom’ mutant, *pyp1-1(H7L)*, defective in xanthophyll esterification (Ariizumi *et al*., 2014). CCS altered the compositions and the flux in the carotenoid pathway of *CCS*-transformed tomato fruit, resulting in higher accumulation of total carotenoids, *β*-carotene, and xanthophyll esters. The results from overexpressing *CCS* in the mutant background indicated that xanthophyll esterification process is important for carotenoid biosynthesis and accumulation. Furthermore, we introduced *CCS* into five different tomato inbreds through controlled crosses, which resulted in remarkable improvements of *β*-carotene levels. CCS-overexpression also increased volatile apocarotenoids derived from β-carotene with no significant impact on plant growth and yield. We demonstrated that the tomato fruit biofortified with high provitamin A and ketocarotenoids would be an excellent dietary source for human nutrition and to address VAD.

## Materials and Methods

### Plant materials and growth conditions

Seeds of wild-type (WT) *Solanum lycopersicum* L. cv Micro-Tom, and *pale yellow petal 1-1* mutant (*pyp1-1*, TOMJPE-H7L, Fig. S1) in ‘Micro-Tom’ background, were from TOMATOMA, Japan (Saito *et al*., 2011). The mutant *pyp1-1* exhibits an inability to synthesize xanthophyll esters (Ariizumi *et al*., 2014). Seeds of LA2374 (‘Caro Red’), LA2377 (‘Purple Calabash’), and LA4044 (*Solanum pennellii*, IL3-2, (Alseekh *et al*., 2013) were from the Tomato Genetics Resource Center (University of California, Davis, CA, U.S.A.). Seeds of FL7907B were from Dr. S.Hutton (GCREC, University of Florida, Wimauma, FL, U.S.A.). Seeds of ‘Jaune Flamme’ were from Tomato Grower’s Supply Company (Fort Myers, FL, U.S.A.). Seedlings were grown in a potting medium (Jolly Gardener, C25) supplied with controlled-release fertilizer (Florikan, N: K:P=18:6:8). Growing conditions were as follows: 16-h-day/8-h-night photoperiod, 25°C/22°C day/night temperature, 50% relative humidity, and ∼150 µmol m^−2^ s^−1^ light intensity supplemented by fluorescent lights.

### cDNA Cloning

Total RNA was isolated from ripe pepper pericarp of RJ107(6)A3 (Maquilan *et al*., 2020) using RNeasy plant mini kit (Qiagen, Germany). First-strand cDNA was synthesized using SuperScript^®^III First-Strand Synthesis System (Invitrogen, U.S.A.). Capsanthin/capsorubin synthase (CCS, Capana06g000615, PP418871.1) protein coding sequence (CDS) minus stop codon were amplified from first strand cDNA using specific primers listed in Table S1. The gel-purified amplicons were cloned into an entry vector pENTR^™^/D-TOPO^®^ (Invitrogen, U.S.A.). *CaCCS* cDNA from the entry vector were subcloned into plant expression vector pK7FWG2.0 by Gateway LR recombination reaction (Invitrogen, U.S.A.) to generate pK7FWG2.0-CaCCS-eGFP. Following all cloning operations, sequence verification was done by Sanger sequencing.

### Virus-Induced *ccs*-Silencing in Pepper Fruit

The *CCS* knock-down pepper lines were generated by the VIGS modified from (Liu *et al*., 2002). The target region for silencing was designed by the SGN VIGS Tool (https://vigs.solgenomics.net). The output suggested the best target region and no potential off-target in that 460 bp fragment of *CaCCS*. It was amplified from pepper fruit cDNA using specific primers listed in Table S1. After subcloning into pTRV2-MCS, which was obtained along with pTRV1 from the Arabidopsis Biological Resource Center, pTRV2-CCS was generated. This plasmid was transformed into *Agrobacterium tumefaciens* strain GV3101. The agroinfiltration of pepper plants was performed as previously described (Kim *et al*., 2017). *Agrobacterium* strains GV3101 harboring pTRV2-CCS and pTRV1 were mixed 1:1 and infiltrated onto open cotyledons using 1-mL syringe without needles. The *Agrobacterium* strain GV3101 containing pTRV2-MCS was used as a control. Infiltrated plants were kept under dark and high humidity conditions overnight, followed by normal conditions.

### Quantitative Real-Time PCR

RNA extraction and single-stranded cDNA were prepared as described in ‘cDNA Cloning’. The expression level of *CCS* gene was estimated by quantitative RT-PCR using green fast qPCR blue mix (AzuraView^TM^, MA, U.S.A.) with gene-specific primers (Table S1). The relative expression levels of targeted genes were quantified using the equation of 2^^–ΔCT^, normalized to the gene encoding ubiquitin-conjugating enzyme E2 (Cheng *et al*., 2017). Each qPCR reaction was run in four biological and three technical replicates.

### Plasmid Construction and Carotenoid Production in *E. coli*

Background plasmids, pAC-LYCipi (plasmid # 53279) and pAC-VIOL (Plasmid #53087), were obtained from Addgene, which was developed previously to produce lycopene and violaxanthin, respectively, in *Escherichia coli* (Cunningham *et al*., 2007; Neuman *et al*., 2014). The background plasmid was transformed into an *E. coli* strain, SixPack, derived from BL21(DE3) (Lipinszki *et al*., 2018). Pepper and tomato cDNAs were prepared as described in ‘cDNA Cloning’. The cDNA of CaCCS, SlLYCB1 (Solyc04g040190), and SlLCYB2 (Solyc10g079480) minus transient peptide sequence was amplified using specific primers listed in Tab. S1. After subcloning into pET30a (Novagen), pET30-CaCCS, pET30-SlLCYB1 and pET30-SlLCYB2 were generated, sequenced, and used for *E. coli* transformation.

pET30a plasmids, including empty vector (pET30a), pET30-CaCCS, pET30-SlLCYB1, and pET30-SlLCYB2, were transformed into chemically competent SixPack cells harboring the background plasmid encoding carotenoid biosynthetic genes (pAC-LYCipi or pAC-VIOL).

Three individual colonies for each transformation were picked and validated through colony PCR. After validation, they were inoculated separately in 10-mL LB liquid media containing 30 mg L^−1^ of chloramphenicol and 50 mg L^−1^ of kanamycin. The starter cultures were grown at 37 °C until saturation. In the following day, 1 ml of overnight culture was used to inoculate 100 ml of Terrific Broth (TB) medium containing 30 mg L^−1^ of chloramphenicol and 50 mg L^−1^ of kanamycin in a 250-ml Erlenmeyer flask. The cultures were grown at 200 rpm at 37 °C until an absorbance at 600 nm of 1.0 was achieved. The cells were chilled on ice for 10 min and induced with a final concentration of 0.1 mM isopropyl *β*-D-1-thiogalactopyranoside (IPTG). The cultures were returned to the shaker and grown for 24 h at 200 rpm at 16°C in the darkness. Then, the cultures were shaken at 25°C for an additional 60 h at 200 rpm in the darkness. The cells were harvested by high-speed centrifugation, which were used for the extraction and chromatographic analysis of carotenoids.

### Plant transformation

*Agrobacterium*-mediated transformation of tomato was performed using the method of Sun et al., (2006) with modifications. *Agrobacterium* GV3101 harboring pK7FWG2.0-CaCCS-eGFP were grown in YEP to prepare *Agrobacterium* inoculum. Surface-sterilized seeds of WT ‘Micro-Tom’ and *pyp1-1* mutant were germinated on half-strength MS medium (Murashige & Skoog, 1962). Cotyledon explants from 10-day old seedlings were placed on a preculture medium for 48h, blotted dry and placed on a co-cultivation medium at 25°C in the dark for 48h. Subsequently, the explants were transferred to the callus induction medium for two weeks and then to shoot induction medium until shoots formed. Elongated shoots were excised and transferred to the rooting medium. Rooted shoots were transplanted to soil. Media components are listed in Table S2. Genomic PCR with T-DNA-specific primers combining flower phenotyping was performed to identify the putative transformants. More than a hundred of T_0_ plants were screened (257 for WTCCS and 163 for CCS). Validated T_0_ transgenic lines were grown to collect seeds, from which representative WTCCS transgenic lines and CCS transgenic lines at T_2_ generation were used for further analyses.

The transgene copy numbers were evaluated using qPCR and serial dilution of genomic DNA against systemin and phytoene desaturase genes in the genome as the endogenous references (Chetty *et al*., 2013; Kanwar *et al*., 2022). Data on representative transgenic lines presented in Figures 2 to 8 have been verified to contain a single copy of CCS integrated into their genomes. Data on additional transgenic lines are presented in Figures S8 and S9.

### Control– and *CCS*-derived tomato lines

Five tomato inbreds, ‘FL7907B’, ‘LA2377’, ‘LA2374’, ‘Jaune Flamme’, and ‘LA4044’, were selected as maternal lines. The transgenic lines, WTCCS46 and WTCCS59 generated in this study and ‘Micro-Tom’ (WT background) were used as paternal lines. Pollen from the WTCCS46 or WTCCS59 was transferred onto the stigma of emasculated flowers of the selected lines, to generate *CCS*-derived lines. Similar crosses were made between WT ‘Micro-Tom’ and the selected lines to generate control-derived lines. Seeds of ripe fruit resulting from crosses were collected and germinated to generate the F1 plants. Ten plants from each cross were grown as biological replicates. All F1 plants harboring the transgene were validated by genomic PCR with T-DNA-specific primers (Tab. S1). The F1 plants were grown in a temperature-controlled greenhouse in 5-gallon containers filled with potting medium and fertilized with the Tomato Plant Food (MiracleGro, Marysville, OH, U.S.A.) every week. The ripe fruit from all lines were collected for analyses.

### Carotenoid analysis

Carotenoids were extracted from pepper or tomato pericarp, or *E coli* cells following the method of Wahyuni (2011). The pooled organic solvent layers from extracts were combined, dried under a nitrogen stream, and were used either for direct quantification (unsaponified) or for saponification based on the method of Mínguez-Mosquera & Hornero-Méndez (1993). Quantification of carotenoids was performed by HPLC-UV/Vis following the method adapted from Delgado-Pelayo & Hornero-Méndez (2012). The chromatographic system comprised an Agilent 1100 chromatograph connected to an Agilent G1314A UV/Vis detector operated by ChemStation software. The chromatographic conditions are listed in Table S3. Lycopene and *β*-carotene standards were from Sigma-Aldrich (MO, U.S.A.). Zeaxanthin, lutein, and zeaxanthin dipalmitate were from Cayman. Capsanthin and capsorubin were purified from pepper fruit by column chromatography using the methods of Cvetković & Marković (2008). Each carotenoid concentration was calculated using the standard curves. The xanthophyll ester fractions were quantified using the calibration curve of zeaxanthin dipalmitate since standards for most other xanthophyll esters were not commercially accessible. Concentration was calculated as µg per gram of fresh weight (µg g^−1^ f.wt.).

### Analysis of flavor volatiles from ripe fruit

Freshly harvested ripe fruit from greenhouse-grown control and *CCS*-derived hybrids were used to quantify volatile apocarotenoids (Tieman *et al*., 2006). Briefly, a 100 g sample of chopped fruit was placed inside a glass tube through which air filtered through a hydrocarbon trap (Agilent, Palo Alto, CA) was flowing through for 1 h. The volatile apocarotenoids were collected on a Super Q column, which was then eluted with methylene chloride after the addition of nonyl acetate as an internal standard. VOC were separated on an Agilent (Palo Alto, CA) DB-5 column and analyzed using an Agilent 6890N gas chromatograph. Compounds were identified based on retention times compared to standard compounds (Sigma Aldrich, St Louis, MO). The volatile amounts were computed based on peak areas calibrated by internal standards and expressed as ng^-1^ gfwt^-1^ h^-1^.

## Results

### Characterization of capsanthin/capsorubin synthase (CCS) in pepper carotenoid pathway

A Pearson correlation analysis on the relationship between the contents of paired carotenoids from 32 pepper varieties adopted from a previous study (Wahyuni *et al*., 2011) showed that capsanthin and capsorubin contents were positively correlated with *β*-carotene contents (r=0.88) (Fig. S2), suggesting a role for CCS in accumulation of capsanthin, capsorubin, and *β*-carotene in pepper fruit. To confirm CCS’ function in the carotenoid pathway, *CCS* expression in pepper fruit was silenced through virus-induced gene silencing (VIGS). The relative expression of the *CCS* transcript dropped >10-fold in *ccs*-silenced fruit compared to the mock, paralleling with the yellow color of ripe fruit as opposed to the red of the mock control (Fig. 1a). The carotenoid profile of *ccs*-silenced fruit showed a distinguishable difference from that of mock-treated fruit (Fig. 1b). Compared to mock-treated fruit, total carotenoids were significantly reduced in the *ccs*-silenced fruit, among which xanthophyll esters and carotenes decreased by 61.5% and 37.4% respectively (Fig. 1c, f). Specifically, lutein and *ɑ*-carotene contents in the *ɑ*-branch of the carotenoid pathway were elevated by 13.2 and 1.6-fold respectively when *CCS* expression was silenced (Fig. 1d, f). Oppositely, capsanthin and capsorubin levels in the *β*-branch were reduced by 96% and 99% respectively in *ccs*-silenced fruit (Fig. 1f). The silencing of the *CCS* gene in pepper fruit reduced not only capsanthin and capsorubin, but also *β*-carotene levels by 46% in pepper fruit (Fig. 1e, f), consistent with the results of correlation analysis. In the *β*-branch, the amounts of other carotenoids, except for (*all E*) violaxanthin and (*9’Z*) violaxanthin, were reduced significantly as well in *ccs*-silenced fruit (Fig. 1e, f).

**Figure 1.**
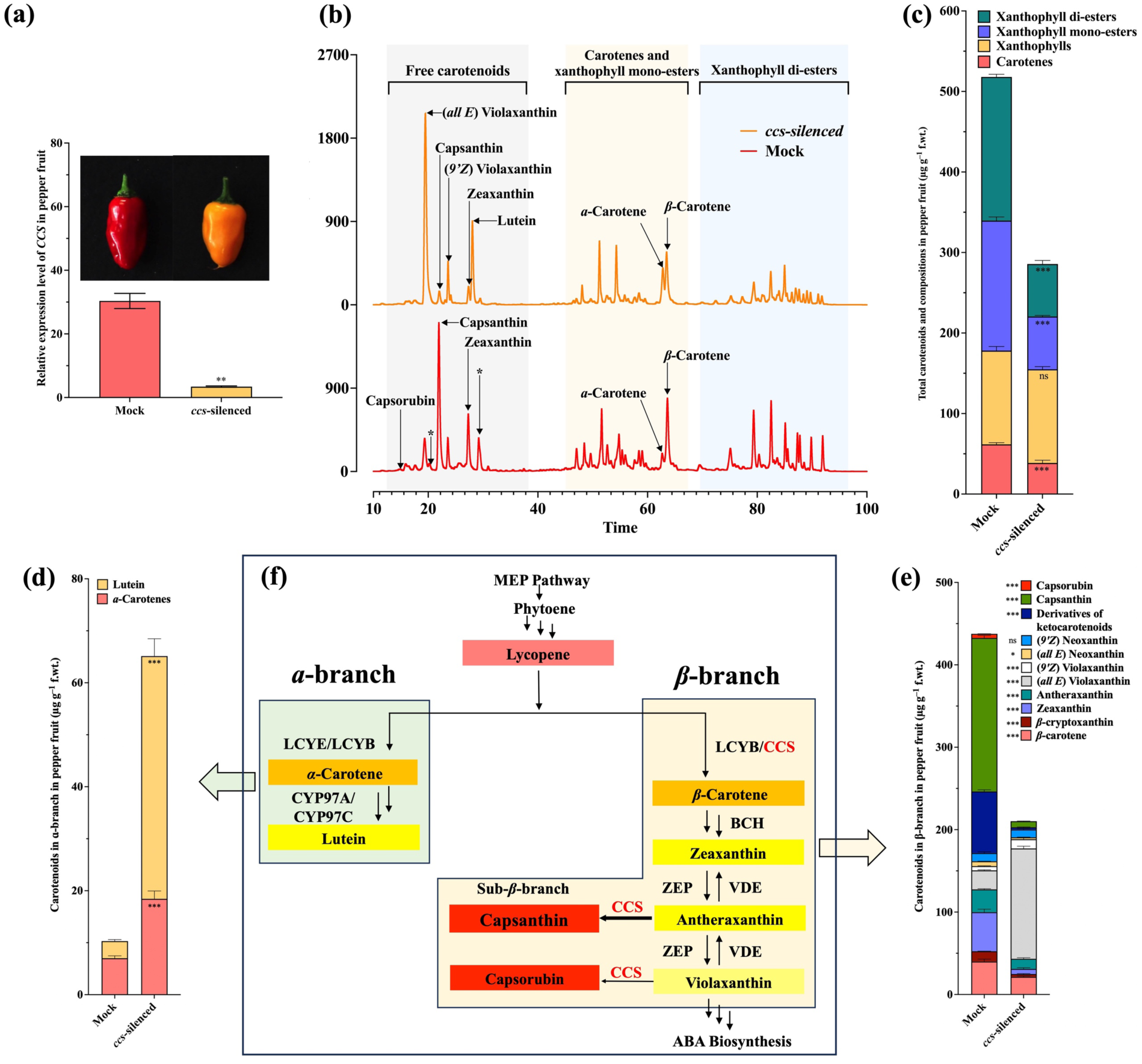
Characterizing the role of capsanthin/capsorubin synthase (CCS) in pepper. (a) The phenotype of *ccs*-silenced pepper fruit and relative expression level of *CCS* gene in *ccs*-silenced pepper fruits compared to mock-treated. Mock treatment refers to the pepper plants infected with VIGS-empty vector without harboring the gene sequence of interest (*CaCCS*). Data of quantitative RT-qPCR are presented as mean ± standard deviation of four biological replicates and three technical replicates. An unpaired t-test was performed (*: P<0.05; **: P <0.01). (b) HPLC chromatograms of mock or *ccs*-silenced pepper fruit carotenoids through the VIGS system. HPLC analyses were performed without saponification. ‘*’ indicated the derivatives of capsanthin and capsorubin (Kim *et al*., 2009). (c) Total carotenoids and their compositions in pepper fruit. HPLC analyses were performed without saponification. (d–e) Carotenoid compositions in the mock or *ccs*-silenced pepper fruits pepper fruit determined after saponification. Data are presented as mean ± standard deviation of four biological replicates. Two-way ANOVA was performed followed by Šídák’s multiple comparisons test (*: P<0.05; **: P<0.01; ***: P<0.001). (f) The carotenoid pathway in plants. The precursor, geranylgeranyl diphosphate, is synthesized from the methylerythritol 4-phosphate (MEP) pathway. Phytoene synthase (PSY) is the first committed step leading to the carotenoid phytoene. Lycopene is synthesized from phytoene after multiple steps, including desaturation and isomerization. Subsequently, lycopene is converted into *ɑ*-carotene or *β*-carotene forming two sub-branches, *ɑ*– and *β*-branches in the following steps, highlighted in green and yellow color backgrounds, respectively. In the *β*-branch, the specific steps in red pepper fruit, produce specialized ketocarotenoids, capsanthin and capsorubin, to generate the sub-β-branch. LCYB, lycopene *β*-cyclase; LCYE, lycopene *ɑ*-cyclase, CYP97A/C, cytochrome P450-type monooxygenase 97A/C; BCH, *β*-carotene hydroxylase; ZEP, zeaxanthin epoxidase; the sub-β-branch indicate the carotenoid flow in pepper fruit.

### Functional validation and comparison of pepper CCS and tomato LCYBs

To compare the enzymatic function of pepper CCS and tomato LCYBs, they were heterologously expressed in an *E. coli* system, respectively (Fig. 2). First, we tested their lycopene cyclase activity in *vivo* (Fig. 2a). The *E. coli* cells accumulated 133.9 µg g^−1^ f.wt. cells of lycopene when it harbored the background plasmid, pAC-LYCipi, and empty vector, pET30a (Fig. 2b&c). When the additional CCS or LCYBs were expressed along with pAC-LYCipi, *β*-carotene was detected in *E. coli* cells (Fig. 2b–d). Specifically, pepper CCS harboring cells accumulated 158.0 µg g^−1^ f.wt. cells of *β*-carotene, accounting for 96% of total carotenoids (Fig. 2b–d). However, both tomato LCYBs harboring cells accumulated 31.7∼37.0 µg g^−1^ f.wt. cells of *β*-carotene and 48.5 and 63.4 µg g^−1^ f.wt. cells of lycopene remaining in *E coli* cells (Fig. 2b–d). Subsequently, to further test the unique function of CCS in ketocarotenoid biosynthesis, another background plasmid, pAC-VIOL, was transformed in *E. coli* to produce precursors, antheraxanthin or violaxanthin (Fig. 2e). The *E. coli* harboring pAC-VIOL along with the empty vector accumulated small amounts of xanthophylls including violaxanthin, antheraxanthin and zeaxanthin (Fig. 2f). However, when CCS or LCYBs were expressed with pAC-VIOL, the total carotenoids were increased (Fig.2f). The substrate, antheraxanthin, for ketocarotenoid synthesis was increased 1.1 to 1.6-fold in LCYBs harboring *E. coli*, while it remained at similar level in CCS harboring *E. coli* (Fig. 2g). Accordingly, capsanthin was detected in CCS harboring *E. coli* but not in LCYBs harboring *E. coli* (Fig. 2h).

**Figure 2.**
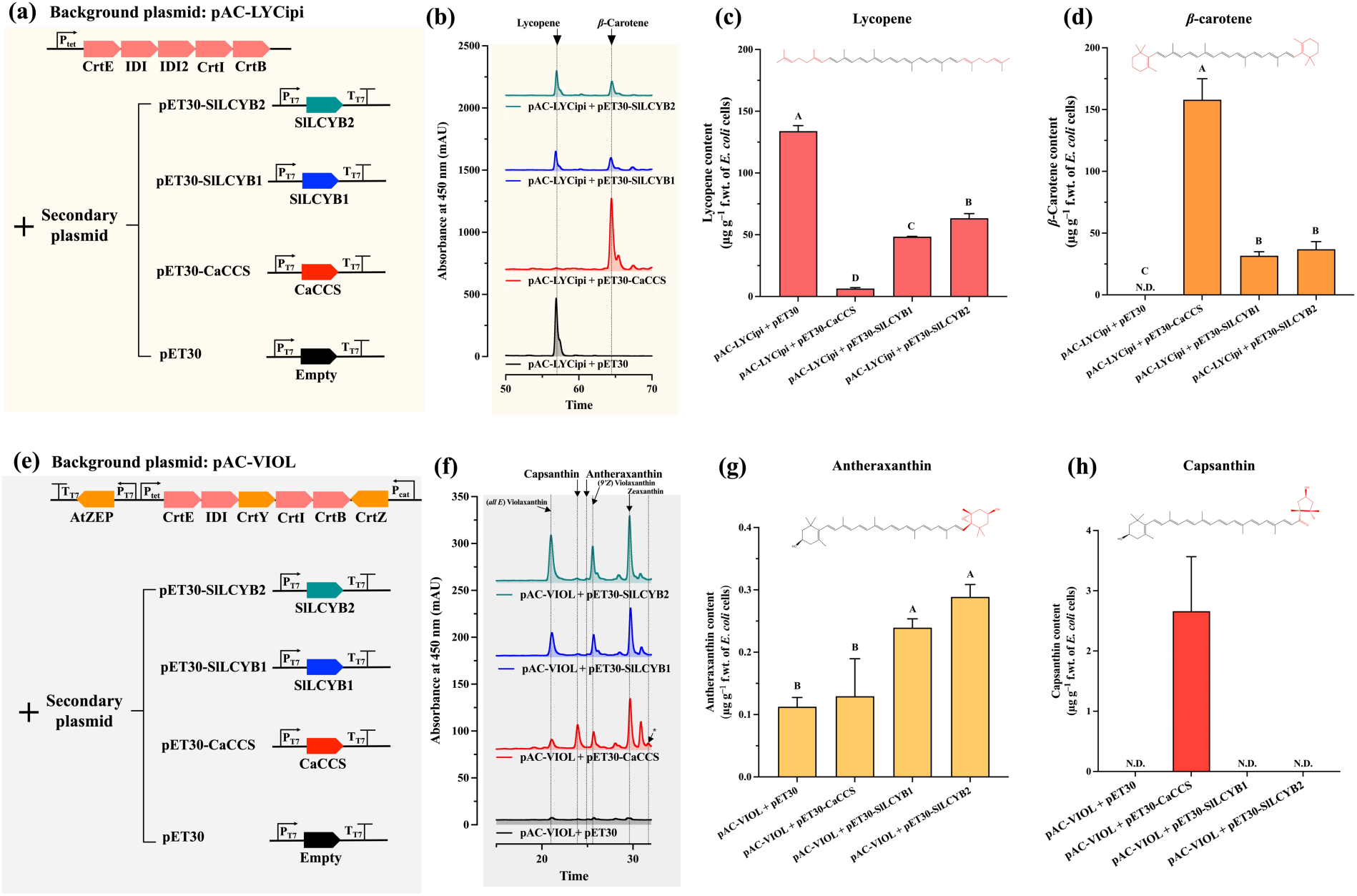
Functional validation and comparison of tomato LCYBs and pepper CCS in *E coli*. a. Schematic representation of different plasmid combinations used for lycopene and *β*-carotene production in *E. coli* cells. The *E. coli* strain, SixPack, harboring the background plasmid, pAC-LYCipi, can produce lycopene. A secondary plasmid containing tomato LCYB1, LCYB2, or pepper CCS was co-transformed with pAC-LYCipi in SixPack. b. The corresponding HPLC chromatograms to the pAC-LYCipi with pET30 (Empty vector), pET30-CaCCS, pET30-SlLCYB1, or pET30-SlLCYB2, respectively. Lycopene (c) and *β*-carotene (d) contents in SixPack harboring different plasmid combinations. e. Schematic representation of different plasmid combinations used for xanthophyll production in *E. coli* cells. The *E. coli* strain, SixPack, harboring another background plasmid, pAC-VIOL, can produce zeaxanthin without IPTG induction and violaxanthin with IPTG induction. Similar as (a), a secondary plasmid containing tomato LCYB1, LCYB2, or pepper CCS was co-transformed with pAC-VIOL in *E. coli*. f. The corresponding HPLC chromatograms to the pAC-VIOL with pET30 (Empty vector), pET30-CaCCS, pET30-SlLCYB1, or pET30-SlLCYB2, respectively. ‘*’ indicated the derivatives of ketocarotenoids. Antheraxanthin (g) and capsanthin (h) contents in SixPack harboring different plasmid combinations. The carotenoid contents were calculated using the peak area of the HPLC chromatogram against standards, which are expressed as µg per gram fresh weight of *E. coli* cells. Data of c–d and g–h are presented as mean ± standard deviation of three biological replicates based on data from HPLC analyses. One-way ANOVA was performed, followed by a mean separation test using the Tukey test and means significantly different at p = 0.05 are shown with different letter codes.

### Phenotypic alteration of *CCS*-overexpression tomato lines

In a control tomato WTEV21 (wildtype background transformed with empty vector, line21) and 1-1EV11 (*pyp1-1(H7L)* background transformed with empty vector, line11), ripe fruit remained red comparable to WT and *pyp1-1(H7L)*, showing no visible differences generated by the empty vector (Fig. 3a, e). When tomato was engineered for constitutive expression of pepper *CCS*, no obvious difference in plant growth and yield was observed, compared to the control lines (Fig. S3). However, the ripe fruit color changed from red to red-orange (WTCCS46 line, a representative line of wildtype background transformed with *CCS*, Fig. 3a). In mutant background, *pyp1-1(H7L)*, the ripe fruit color turned orange (CCS55 line, Fig. 3e). The changes in carotenoid profiles behind the fruit color shift were further investigated through HPLC-UV/Vis analysis (Fig. 3b–d, f–h). The red fruit from the control lines, WT, WTEV21, *pyp1-1(H7L)*, and 1-1EV11 contained predominantly lycopene and tiny amounts of other carotenoids, including *β*-carotene, (*all E*) violaxanthin, and lutein (Fig. 3b–d, f–h). In addition, xanthophyll esters were detected at a low level in the WT and hardly in the *pyp1-1(H7L)* background (Fig. 3d, h). In the ripe fruit of *CCS*-overexpression lines, regardless of WT or *pyp1-1(H7L)* background, capsanthin was identified (Fig. 3b, f). The primary carotenoid of tomato, lycopene, decreased while *β*-carotene, (*all E*) violaxanthin, and xanthophyll esters increased compared to control lines (Fig. 3b–d, f–h).

**Figure 3.**
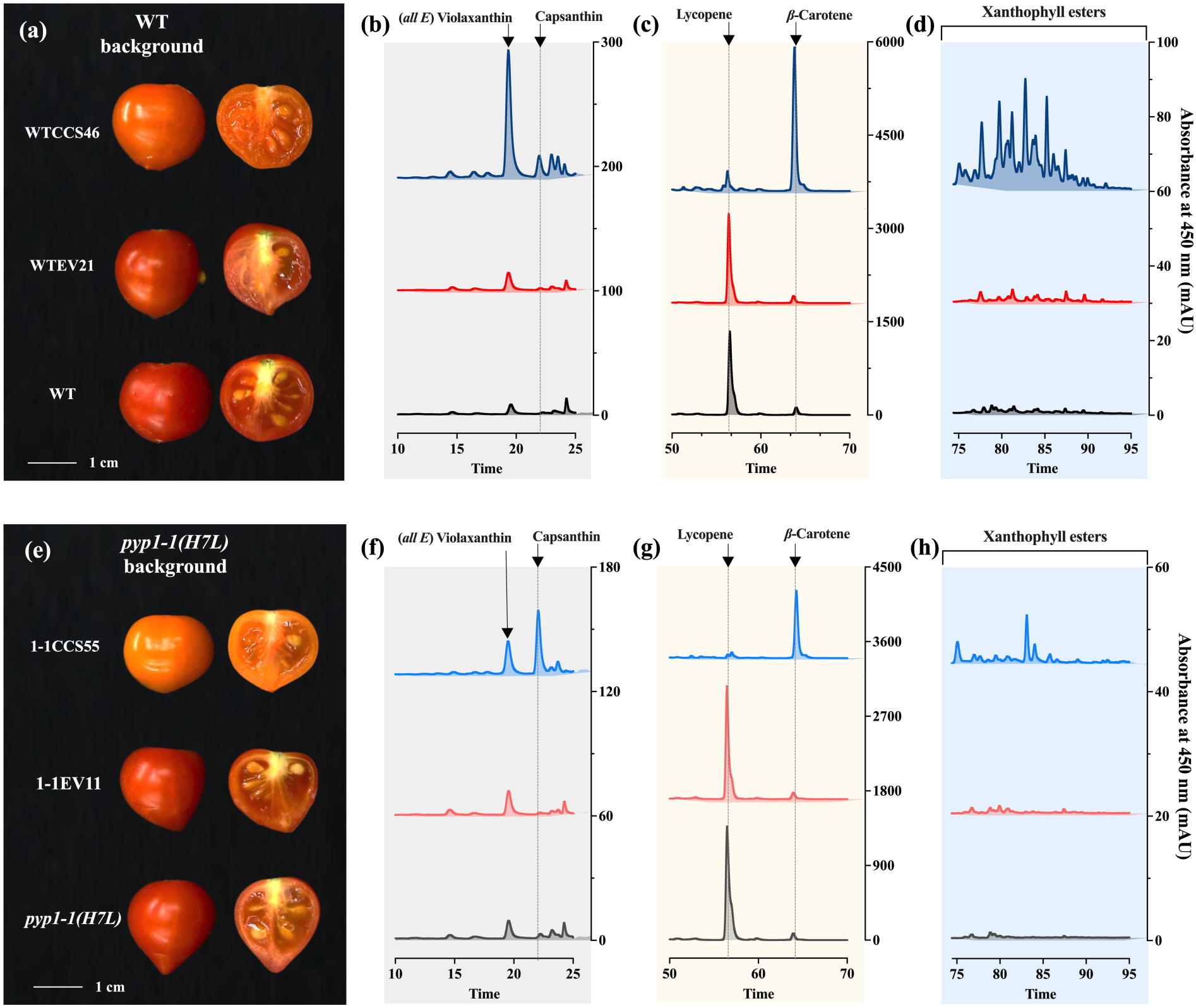
Phenotypes and carotenoid profiles of tomato fruit. Overexpression of pepper *CCS* was performed in ‘Micro-Tom’ background, including two genotypes, wildtype (WT) (a–d) and *pyp1-1(H7L)* mutant for an acyltransferase known in xanthophyll esterification (e–h). (a) The phenotypes of tomato fruits from ‘Micro-Tom’ WT, ‘Micro-Tom’ transformed with empty vector, WTEV21, and a representative line, WTCCS46, from multiple independent *CCS*-transformed lines in ‘Micro-Tom’ WT background. (b–d) The corresponding HPLC chromatograms to ‘Micro-Tom’ WT, WTEV21, and WTCCS46 lines. (e) The phenotypes of tomato fruits from ‘Micro-Tom’ mutant, *pyp1-1(H7L)*, *pyp1-1(H7L)* transformed with empty vector, 1-1EV11, and a representative line, 1-1CCS55, from multiple independent *CCS*-transformed lines in *pyp1-1(H7L)*, background. (f–h) The corresponding HPLC chromatograms to ‘Micro-Tom’ mutant, *pyp1-1(H7L)*, 1-1EV11, and 1-1CCS55.

### Alteration of carotenoid pathway and *β*-carotene enhancement in *CCS*-transformed tomato

Carotenoid profiles in different tomato lines indicated the alteration of carotenoid compositions (Figs.3&4a). Specifically, when total carotenoids in tomato fruit were quantified, the fruit of WTCCS lines were all enhanced compared to WT and WTEV21 lines (Fig. 4b). Their fruit had increments of 33–69% more compared to WT and WTEV21 lines. Nevertheless, total carotenoids in 1-1CCS lines decreased by 20–28% compared to *pyp1-1(H7L)* and 1-1EV11 (Fig. 4b). In addition to total carotenoids, the carotenoid compositions were changed in *CCS*-overexpression lines (Fig. 4c–g). In the red tomato of WT and *pyp1-1(H7L)* backgrounds, the carotenoid fraction was mainly lycopene, which is the final carotenoid synthesized before the pathway was split into *ɑ*-or *β*-branches (Fig. 4c). Specifically, the amounts of lycopene, in WT and WTEV21 lines, decreased from 85% of total carotenoids to 3–9% in WTCCS lines (Fig. 4c). Similarly, in the mutant background, lycopene levels decreased from 87% of total carotenoids to 6–11% in *pyp1-1(H7L)* and 1-1EV11 (Fig. 4c).

**Figure 4.**
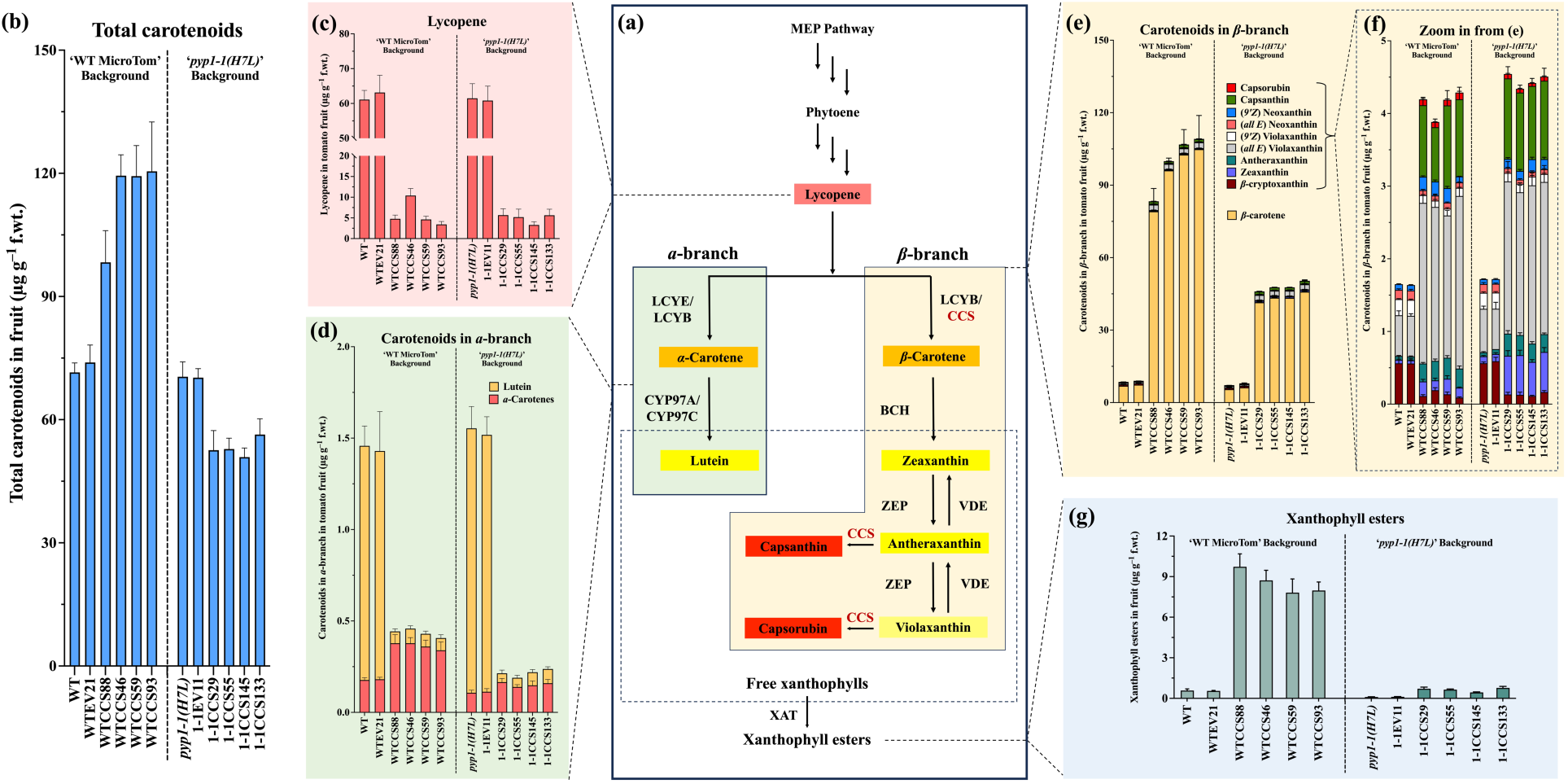
Compositional alterations in the carotenoid profiles of tomato in *CCS*-transformed tomato. (a) The carotenoid pathway in plants is listed in the black box at the center of this figure. Two branches, *ɑ* and *β*-branch, are highlighted in green and yellow backgrounds. The xanthophylls in the dash-line box were available to be esterified under the catalysis of xanthophyll acyltransferase (XAT). Total carotenoids (b) and lycopene contents (c) in the fruit of control and independent *CCS*-transgenic lines in both WT ‘Micro-Tom’ and *pyp1-1(H7L)* backgrounds. (d) Carotenoid compositions, including lutein and *ɑ*-carotene, in the *ɑ*-branch of carotenoid pathway. (e) Carotenoid compositions, including *β*-carotene, *β*-cryptoxanthin, zeaxanthin, antheraxanthin, violaxanthin, neoxanthin, capsanthin and capsorubin in the *β*-branch of carotenoid pathway. (f) A zoom-in view of carotenoids in *β*-branch from (e) except for *β*-carotene. (g) Xanthophyll esters in the fruit of control and independent *CCS*-transgenic lines in both WT ‘Micro-Tom’ and *pyp1-1(H7L)* backgrounds. Data of b–g are presented as mean ± standard deviation of a minimum of four biological replicates based on data from HPLC analyses. One-way ANOVA was performed followed by mean separation test using the Tukey test (*: P<0.05; **: P<0.01; ***: P<0.001), details of which are shown in the supplementary Excel file.

Accordingly, other compositions downstream of the carotenoid pathway were altered when *CCS* was overexpressed in both genetic backgrounds (Fig. 4d–g). Although the amounts of carotenoid in the *ɑ*-branch were much lower than total carotenoids, lycopene, and *β*-carotene, a significant decrease of carotenoid contents in the *ɑ*-branch was observed (Fig. 4d). In *ɑ*-branch, the amounts of *ɑ*-carotene increased by 0.9–1.1-fold than the WT and WTEV21 lines, and 0.2– 0.5-fold than the *pyp1-1(H7L)* and 1-1EV11 lines (Fig. 4d). However, the lutein content decreased by ∼95% in both genetic backgrounds (Fig. 4d). Opposite to the decrease in total amounts in the *ɑ*-branch, most compositions, especially *β*-carotene, in *β*-branch were increased in *CCS*-transformed fruit (Fig. 4e, f). Particularly, fruit *β*-carotene levels enhanced from 7 to 96– 105 µg g^−1^ f.wt. in WTCCS lines (Fig. 4e). Similarly, it increased from 6 to 41–45 µg g^−1^ f.wt. in 1-1CCS lines (Fig. 4e). Compared to WT and WTEV21 lines, about 11–14-fold increases were observed in WTCCS lines (Fig. 4e). Although the increment was lower than the WT background, still about 7–8-fold increases in *β*-carotene levels were observed in 1-1CCS lines compared to *pyp1-1(H7L)* and 1-1EV11 lines (Fig. 4e). The other carotenoids in the *β*-branch also showed significant changes in *CCS*-overexpression lines, although they took minor proportions in this branch (Fig. 4e). Specifically, the amounts of zeaxanthin, antheraxanthin, and (*all E*) violaxanthin were increased by 1.8–3.9, 3.6–4.5, and 2.5–3.2-fold in WTCCS lines than the WT and WTEV21 lines (Fig. 4f). Similar increase patterns as WTCCS lines were observed in 1-1CCS lines. Interestingly, the increase rate of zeaxanthin in 1-1CCS lines was 3.7–4.8-fold, higher than in WTCCS lines (Fig. 4e). Since CCS is not native to tomato, no capsanthin or capsorubin was detected in fruit from non-engineered or empty vector-transformed lines (Fig. 4e). However, the fruit of WTCCS lines contained capsanthin at levels of 0.74–1.14 µg g^−1^ f.wt., which was similar to the amounts in 1-1CCS lines. Capsorubin contents in the fruit of WTCCS lines were 0.07 to 0.1µg g^−1^ f.wt., higher than 0.04–0.06 µg g^−1^ f.wt. accumulating in 1-1CCS lines (Fig. 4e).

Free xanthophylls generated in both *ɑ*– and *β*-branches can be esterified. Ripe fruit of WT and WTEV21 lines accumulated 0.55–0.58 µg g^−1^ f.wt. xanthophyll esters (Fig. 3d, 3g). In WTCCS lines, ripe fruit contained more xanthophyll esters at levels of 8–10 µg g^−1^ f.wt., increasing by 12.5–16.8-fold compared to the WT and WTEV21 lines (Fig. 3d, 3g). Although, the *pyp1-1(H7L)* mutant was reported to produce no xanthophyll esters in flowers (Ariizumi *et al*., 2014), relatively small amounts of esters were detected in fruit at the level of ∼0.1 µg g^−1^ f.wt. (Fig. 3h & 3g). Moreover, the degree of xanthophyll esterification was elevated 3∼6 times when *CCS* was overexpressed in the mutant background (Fig. 3h, 3f).

The Pearson correlation analysis showed that the carotenoid pools in WTCCS and 1-1CCS lines were altered in a similar fashion (Fig. 5). Capsanthin and capsorubin contents were positively correlated with the accumulation of *ɑ*-carotene, *β*-carotene, antheraxanthin, (*all E*) violaxanthin, and xanthophyll esters when CCS was overexpressed in both backgrounds. However, the alteration in total carotenoids by overexpressing *CCS* in the WT background exhibited positive correlations with the amounts of capsanthin, capsorubin, and *β*-carotene, opposite to 1-1CCS lines (Fig. 5).

**Figure 5.**
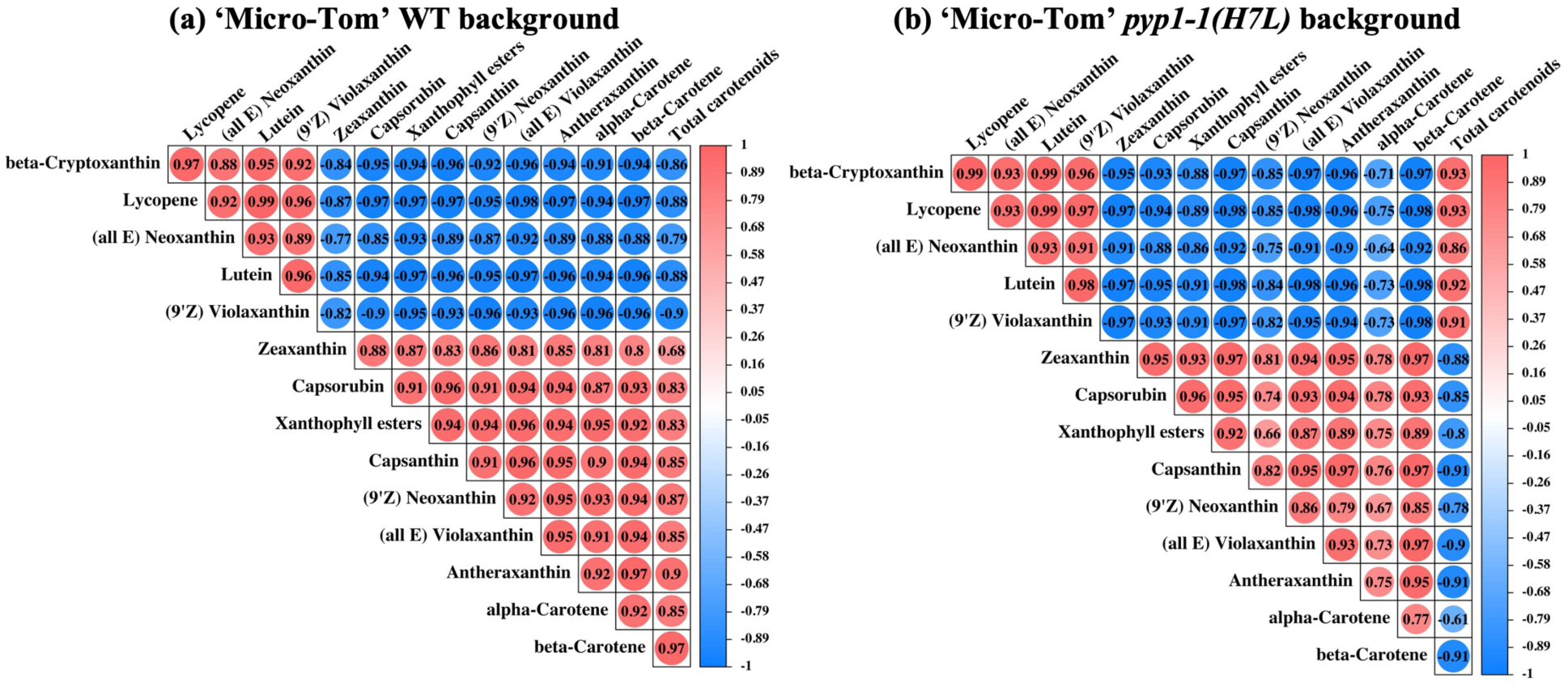
Pearson correlation matrix between each paired carotenoid levels analyzed from ripe tomato fruit. The Pearson correlation coefficient of each paired carotenoid composition is shown in each cell computed based on the carotenoid contents from the set of WT, WTEV and WTCCS lines (a), or *pyp1-1(H7L)*, 1-1EV, and 1-1CCS lines (b) using a R package ‘corrplot’. The Pearson correlation coefficient shown in the cell indicates that it was tested significantly (p-value <0.05), while a tested insignificant correlation would not be shown in the cell. The color intensity and the size of each circle indicated the strength of correlation between each paired carotenoids, in which the more intense the color and larger the circle size are, higher the correlation between each paired carotenoids is. Red indicates positive correlations and blue indicates negative correlations.

### Phenotypic, carotenoid, and apocarotenoid compositional changes in tomato fruit of *CCS*-derived hybrids

To examine CCS function in the fruit of some large-fruited tomato of different genetic backgrounds (Fig. 6a), varieties were selected to do controlled crosses with the transgenic line from WTCCS46 or WTCCS59 lines ‘*CCS*-derived hybrids’ or with WT ‘Micro-Tom’ to generate ‘control-derived hybrids. Ripe fruit color from the selected maternal lines was red for ‘FL7907B’ and ‘LA2377’, orange for ‘Juane Flamme’ and ‘LA2374’, and yellow for ‘LA4044’ (Fig. 6a). The paternal lines WT ‘Micro-Tom’ and WTCCS (line 46) had red and orange ripe fruit color, and different carotenoid compositions, respectively (Fig. 6a).

**Figure 6.**
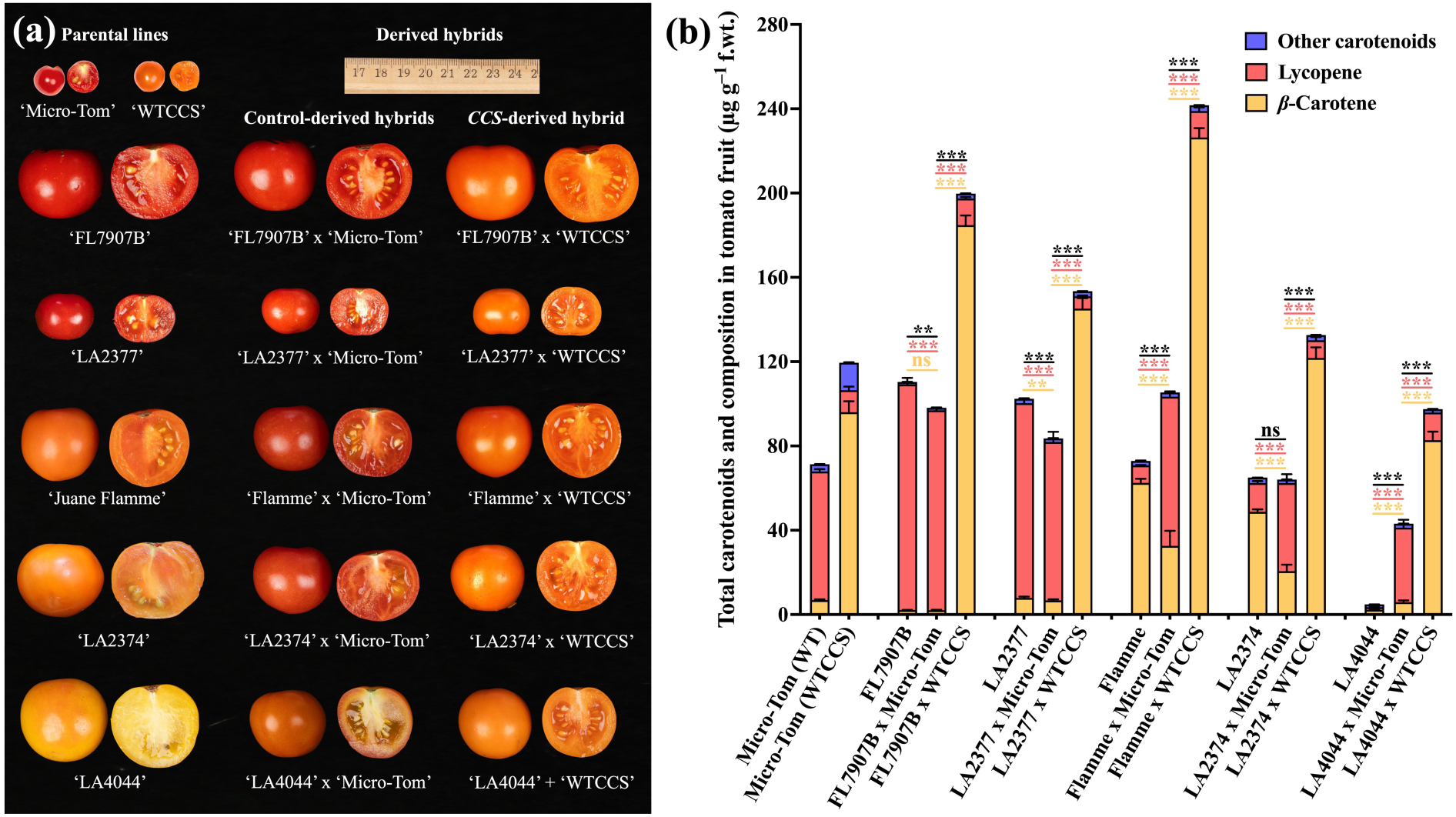
Phenotypic characterization of ripe fruit and carotenoid compositions of parental lines, control– and *CCS*-derived F1 hybrids. (a) Phenotypes of ripe fruit from parental lines, control– and *CCS*-derived hybrids. Five tomato inbred lines, ‘FL7907B’, ‘LA2377’, ‘LA2374’, ‘Jaune Flamme’, and ‘LA4044’, were selected as maternal lines. The transgenic line, WTCCS46, generated in this study and ‘Micro-Tom’ (WT background) were used as paternal lines. (b) Total carotenoids and their compositions in the fruit of parental lines, control– and *CCS*-derived F1 hybrids. Data are presented as mean ± standard deviation of a minimum of four biological replicates based on HPLC analyses. One-way ANOVA was performed followed by mean separation using the Tukey test (*: P<0.05; **: P<0.01; ***: P<0.001). The asterisk color, yellow, pink, and black indicate the mean comparisons of *β*-carotene, lycopene, and total amounts of carotenoids, respectively.

After crossing with WT ‘Micro-Tom’, the fruit of F1 hybrid with ‘Juane Flamme’, ‘LA2374’, and ‘LA4044’ turned red to different degrees while F1 hybrids with ‘FL7907B’ and ‘LA2377’ backgrounds remained unchanged (Fig. 6a). Tomato fruit from F1 hybrids of red tomato varieties, ‘FL7907B’ x ‘Micro-Tom’ and ‘LA2377’ x ‘Micro-Tom’, the total carotenoids in the fruit of control-derived lines were decreased by 11–18%, compared to their corresponding maternal lines (Fig. 6b). When using orange and yellow tomato varieties as maternal parents, fruit from F1 hybrids of ‘Juane Flamme’ x ‘Micro-Tom’ and ‘LA4044’ x ‘Micro-Tom’ showed 0.4 and 7.9-fold increase than their corresponding maternal lines respectively (Fig. 6b). No difference was observed between fruits of ‘LA2374’ and ‘LA2374’ x ‘Micro-Tom’ (Fig. 6b). Specifically, the lycopene contents of control-derived hybrids were increased by 2, 7.5 and 28-fold for ‘Juane Flamme’ x ‘Micro-Tom’, ‘LA2374’ x ‘Micro-Tom’, and ‘LA4044’ x ‘Micro-Tom’, but decreased by 0.11 and 0.19-fold for ‘FL7907B’ x ‘Micro-Tom’ and ‘LA2377’ x ‘Micro-Tom’ respectively, compared to their corresponding maternal lines (Fig. 6b). Besides, except for ‘LA4044’, the *β*-carotene amounts of control-derived hybrids were all decreased by 3– 57%, compared to their corresponding maternal lines (Fig. 6b).

When *CCS* was introduced into the genome of those tomato varieties, all fruit of *CCS*-derived hybrids from these crosses showed color changing to orange, although their color intensity varied depending on their maternal parents (Fig. 6a). Furthermore, the carotenoid compositions were changed thoroughly compared to control-derived hybrids (Fig. 6b). The total carotenoids of the fruits from *CCS*-derived hybrids were increased by 0.8–1.3-fold, compared to their corresponding control-derived hybrids (Fig. 6a). Specifically, lycopene amounts were significantly reduced by 63–92%, compared to their corresponding control-derived hybrids (Fig. 6b). The fold changes of *β*-carotene contents in the fruit from *CCS*-derived hybrids varied depending on their maternal lines (Fig. 6b). Similar to Micro-Tom transformed with *CCS*, *β*-carotene contents in the F1 hybrid tomato fruit generated from red tomato maternal varieties, ‘FL7907B’ x ‘WTCCS’ and ‘LA2377’ x ‘WTCCS’, were increased from 2.1 and 6.7 to 185 and 145 µg g^−1^ f.wt. respectively, (Fig. 6b). In tomato fruit from F1 hybrids generated from orange tomato maternal varieties, ‘Flamme’ x ‘WTCCS’ and ‘LA2374’ x ‘WTCCS’, *β*-carotene contents were 226 and 122 µg g^−1^ f.wt., 2.6 and 1.5-fold more than their corresponding control-derived hybrids (Fig. 6b). F1 hybrid fruit, ‘LA4044’ x ‘WTCCS’, accumulated 5.9 µg g^−1^ f.wt. *β*-carotene, 35 times as in its control-derived hybrid (Fig. 6b). Except for ‘LA4044’, four other CCS-derived hybrids showed 0.2–1.4-fold more *β*-carotene accumulating in fruits than their paternal line, WTCCS46 (Fig. 6b).

In addition to changes in carotenes, capsanthin was detected at levels of 0.22–0.35 µg g^−1^ f.wt. in the fruit of all CCS-derived hybrids, which were lower than the level of 0.74 µg g^−1^ f.wt. in the fruit of the paternal line, WTCCS46 (Fig. 7a). Fruit of ‘Flamme’ x ‘WTCCS’ contained the highest capsanthin content among all five F1 hybrids (Fig. 7a). In all fruit from of F1 *CCS*-derived hybrids, the amounts of xanthophyll esters were significantly increased by 0.13–2.5-fold and 1.3–2.7-fold compared to their corresponding maternal and control-derived hybrids respectively (Fig. 7b). Fruit from ‘Flamme’ x ‘WTCCS’ contained the highest xanthophyll ester content among all five F1 hybrids (Fig. 7b).

**Figure 7.**
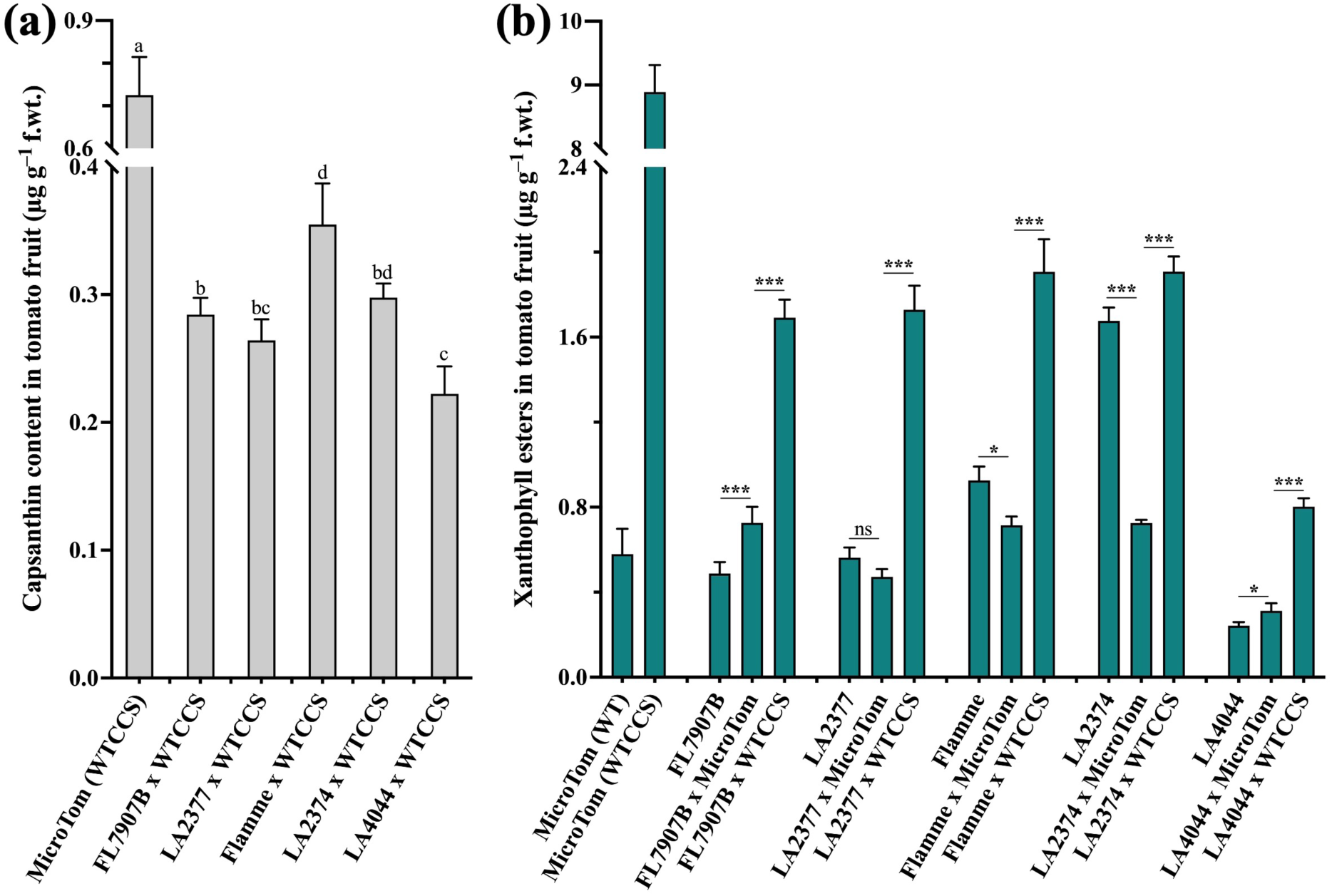
Contents of capsanthin (a) and xanthophyll esters (b) in the fruit of control– and *CCS*-derived hybrids. Data are presented as mean ± standard deviation of a minimum of four biological replicates based on HPLC analyses. One-way ANOVA was performed followed by mean separations using the Tukey test (*: P<0.05; **: P<0.01; ***: P<0.001).

To test whether the volatile apocarotenoids derived from carotenoids, especially those from lycopene and *β*-carotene, changed in *CCS*-derived hybrids, ripe fruits from control and CCS-derived hybrids were collected and analyzed. In control-derived hybrids, ‘Flamme’ x ‘Micro-Tom’ and ‘LA2374’ x ‘Micro-Tom’ with orange fruit as one of the original parents showed higher amounts of *β*-ionone and *β*-cyclocitral, compared to the other control-derived hybrids (Fig. 8a&b). When *CCS* was overexpressed, both *β*-ionone and *β*-cyclocitral amounts were increased by 1.3 to 13.0-fold and 2.7 to 19.1-fold in all five *CCS*-derived hybrids (Fig. 8a&b). The amounts of geranial, geranyl acetone, 6-methyl-5-hepten-2-one, and 6-methyl-5-hepten-2-ol showed a decreasing trend in *CCS*-derived hybrids with one exception for ‘LA2377’ x ‘WTCCS’ and the decrease in geranyl acetone was statistically significant (Fig. 8c–d).

**Figure 8.**
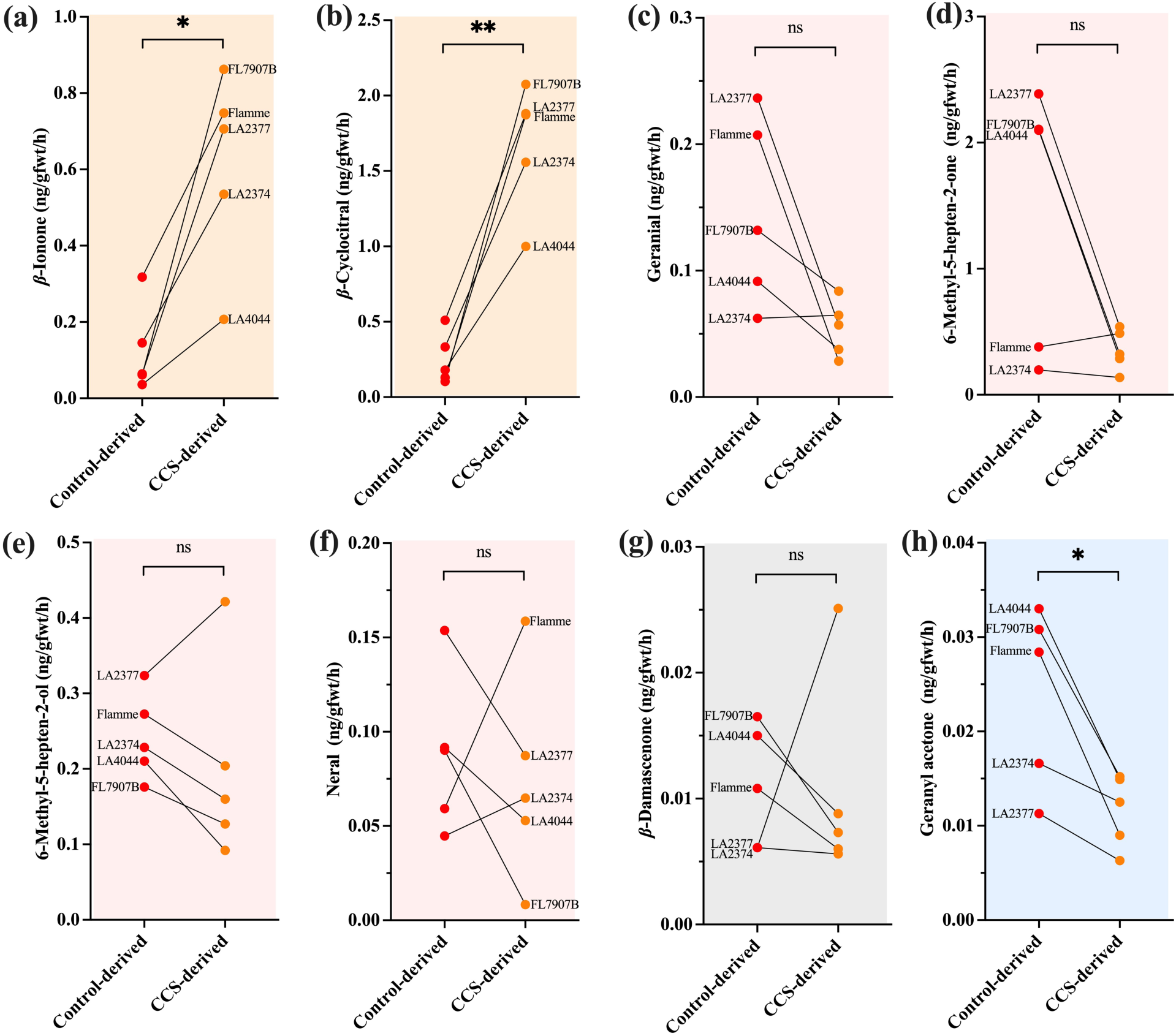
Volatile apocarotenoid derived from carotenoids in ripe tomato from control and CCS-derived hybrids. Beta-ionone (a) and *β*-cyclocitral (b) are derived from *β*-carotene, highlighted in orange background. Geranial (c), 6-methyl 5-hepten-2-one (d), 6-methyl 5-hepten-2-ol (e), and neral (f) are derived from lycopene, highlighted in red background. Beta-damascenone (g) and geranyl acetone (h) are derived from neoxanthin and phytoene respectively, highlighted in gray and blue backgrounds respectively. Data are from 100 g fruit for each plant analyzed by individual runs of GC-MS. ‘FL7907B’, ‘LA2377’, ‘Flamme’, ‘LA2374’, and ‘LA4044’ represent the maternal lines crossed with WT ‘Micro-Tom’ (red dots) or ‘WTCCS’ (orange dots). Asterisks indicate significant (*, p<0.05) or highly significant (**, p<0.01) differences following a paired T-test between the control-derived hybrid group and *CCS*-derived hybrid group (n=5). NS – No significant difference.

### Retinol activity equivalents in engineered fruit

To quantify the nutritional improvement of tomato via *CCS*-overexpression, we evaluated retinol activity equivalents (RAE) computed based on bioactivities of vitamin A and provitamin A (*β*-carotene, *β*-cryptoxanthin, and *ɑ*-carotene) of ripe fruit pericarp from control and *CCS*-overexpression lines (‘Micro-Tom’ background), as well as from the derived hybrids. No difference in RAE was observed between the fruit of WT and WTEV21 (Fig. 9). Nevertheless, fruit from WTCCS lines contained 662–875 µg retinol activity equivalents per 100g f.wt. serving, respectively (Fig. 9). The RAE amounts in fruit from *CCS*-overexpression lines increased by 9.2–13.5 times compared to WT and WTEV21 lines (Fig. 9). However, the RAE amounts enhanced from 48.5–55.0 to 347–384 µg per 100g f.wt. when *CCS* was overexpressed in the *pyp1-1(H7L)* background (Fig. 9). Importantly, the RAE amounts from the fruit of most *CCS*-derived F1 hybrids were observed to be significantly higher levels than transgenic ‘Micro-Tom’ overexpressing CCS (WTCCS46, Fig. 9). Specifically, the fruit of *CCS*-derived F1 hybrids, ‘FL7907B’ x ‘WTCCS’, ‘LA2377’ x ‘WTCCS’, ‘Juane Flamme’ x ‘WTCCS’, ‘LA2374’ x ‘WTCCS’, and ‘LA4044’ x ‘WTCCS’, contained RAE at levels of 1540, 1209, 1885, 1014, and 689 µg per 100g f.wt. serving, respectively, compared to their corresponding control-derived hybrids at levels of 17, 56, 272, 172, and 49 µg per 100 g f.wt. (Fig. 9).

**Figure 9.**
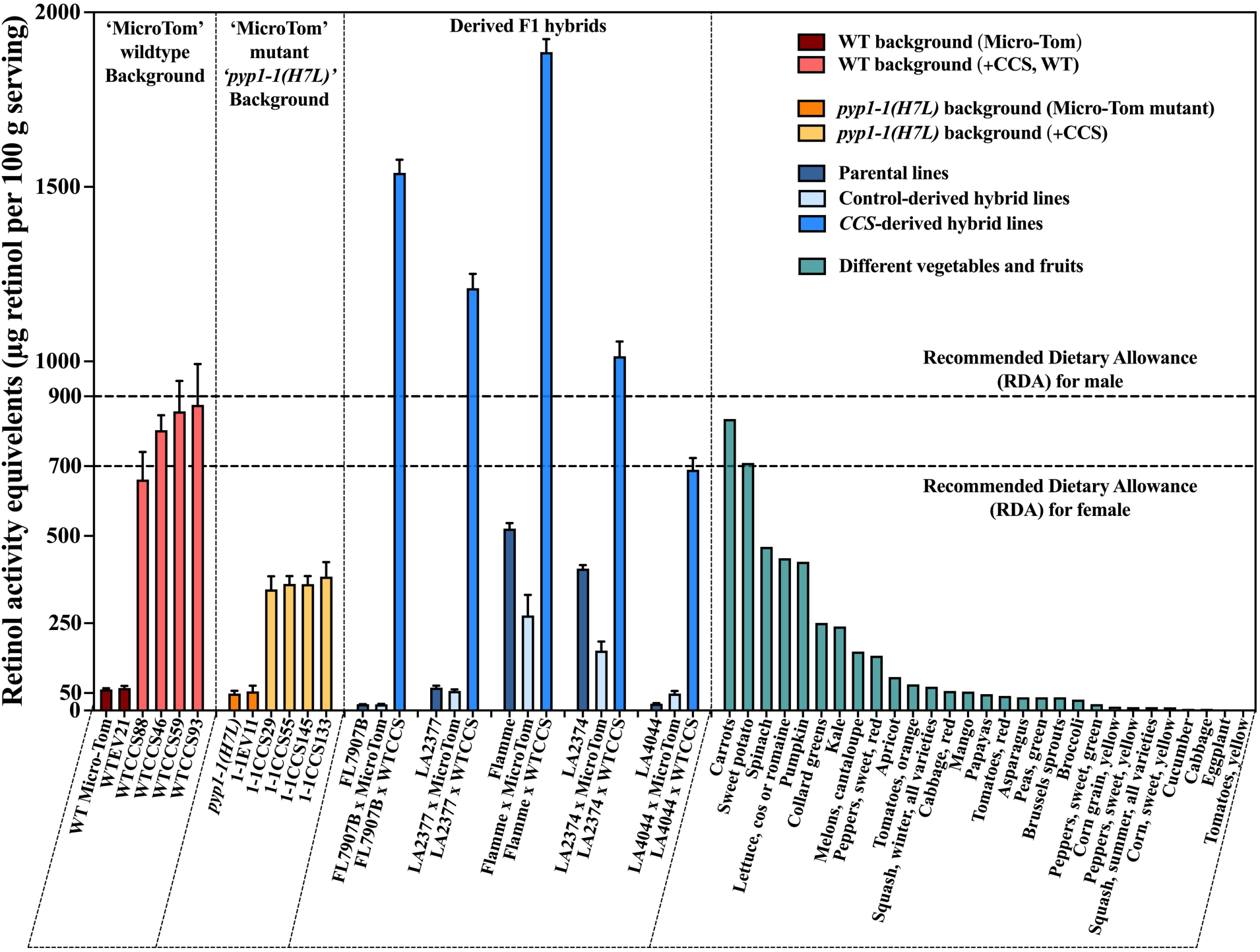
Retinol activity equivalents (RAE) in different vegetables and fruits. RAE values were computed based on the contents of provitamin A, including *β*-carotene, *β*-cryptoxanthin, and *ɑ*-carotene in tomato fruit of engineered lines in this study. The RAE values from different vegetables and fruits, including common sources, were grouped into four, distinguished by dash lines in the figure. The first group depicts data for engineered tomato lines with ‘Micro-Tom’ WT background. The second group contains engineered tomato lines under the ‘Micro-Tom’ *pyp1-1(H7L)* mutant background. The third group contains the parental lines and their derived F1 hybrids. The parental lines included WT ‘Micro-Tom’ and ‘WTCCS Micro-Tom’ (paternal) and five selected inbreds (maternal) ‘FL7907B’, ‘LA2377’, ‘Juane Flamme’, ‘LA2374’, and ‘LA4044’. The fourth group referred to the RAE values of different vegetables and fruits collected from the USDA database (standard reference legacy), whose details are listed in a supplementary Excel file.

## Discussion

Silencing of *CCS* in ripe pepper fruit was conducted to test its role in the carotenoid pathway, which has not been investigated until this study (Fig. 1). The capsanthin and capsorubin formation in fruit was abolished when CCS overexpression was knocked down, with a concomitant change in pericarp color to yellow (Fig. 1a), resembling naturally occurring orange and yellow peppers (Ha *et al*., 2007; Guzman *et al*., 2010). In pepper fruit, CCS draws the carotenoid pathway to form a new sub-branch to produce ketocarotenoids, capsanthin and capsorubin (Fig. 1e, f). When this sub-*β*-branch was abolished by VIGS, the flux went downstream of main *β*-branch to accumulate abundant amounts of violaxanthin instead (Fig. 1b, e). However, the flux was not retained in the main *β*-branch when *CCS* was silenced. In addition to its function in capsanthin and capsorubin biosynthesis, CCS was reported to have LCYB activity *in vitro* (Bouvier *et al*., 1997; Mialoundama *et al*., 2010). Consistent with this, silencing of *CCS* reduced, both cyclization of lycopene and formation of ketocarotenoids. Thus, the limited flow into the *β*-branch caused by the VIGS redirected the flux toward the *ɑ*-branch, to synthesize *ɑ*-carotene and lutein (Fig. 1d).

We observed that the amounts of total carotenoids were reduced remarkably in *ccs*-silenced fruit, suggesting that *CCS* also assisted in the accumulation of total carotenoids, with more ketocarotenoids produced when CCS functioned normally (Fig. 1c, e). Both ketocarotenoids might be preferable substrates for esterification than lutein and violaxanthin in pepper (Fig. 1c, d, f). This was supported by our results that the reduction of capsanthin, capsorubin, and their derivatives was paralleled by an increase of lutein and violaxanthin in *ccs*-silenced fruit, while the esterification of lutein and violaxanthin did not complement the amounts of xanthophyll esters (Fig. 3c), reported storage forms of the carotenoid pool (Li *et al*., 2023). Thus, lack of xanthophyll esterification might cause a flux reduction of the whole carotenoid pathway. Overall, our results elucidated for the first time the critical role of CCS in the metabolic flux and pool of the carotenoid pathway in pepper fruit, although the silencing effect of *CCS* gene through VIGS has been tested in pepper previously (Fig. 1, Tian et al., 2015).

Previous research indicated that pepper LCYB had lycopene cyclization activity and pepper CCS had catalyzed both capsanthin synthesis and lycopene cyclization *in vitro* (Bouvier *et al*., 1997; Mialoundama *et al*., 2010), though these two proteins were not compared *in vivo*. Using an *E. coli* expression platform engineered to produce up to lycopene in the carotenoid synthetic pathway (Fig. 2a) or a strain that could produce xanthophylls (Fig. 2e), we compared pepper CCS with the two tomato LYCBs. Under the same conditions, the *E. coli* strain harboring pepper CCS showed a significantly higher ability to convert lycopene into *β*-carotene compared to the strain containing either LCYB1 or LCYB2 (Fig. 2b–d). Our results are consistent with CCS having a lower K_m_ to lycopene substrate compared to pepper LCYB in a previous study (Mialoundama *et al*., 2010). Accordingly, lycopene in the CCS-harbored strain was depleted by CCS, and much more *β*-carotene was synthesized (Fig. 2d). In another scenario of capsanthin biosynthesis (Fig. 2e), the *E. coli* strain containing CCS with pAC-VIOL plasmid produced capsanthin successfully, while the expression of either LCYB1 or LCYB2 did not result in capsanthin accumulation (Fig. 2f-h). These results proved that tomato LCYBs could not synthesize ketocarotenoid, but CCS did function in capsanthin production in *E. coli*. Particularly, pepper CCS and tomato LCYBs could increase the carotenoid flux downstream to produce more carotenoids, including *β*-carotenes and xanthophylls, indicating that lycopene cyclization might be a limited step in the carotenoid biosynthetic pathway, agreeing with previous studies (Moreno *et al*., 2020; Kössler *et al*., 2021). These results suggested that CCS is superior for *in vivo* modification of carotenoid pools by taking advantage of its multifunctional activities.

Given that CCS could change the flux in the carotenoid pathway based on its multiple functions in pepper fruit and in engineered *E. coli*, and has not been used for directed modification of carotenoid profiles of a ripening fruit yet, its overexpression in the fleshy fruit of tomato allowed us to understand the sink strength in the carotenoid pathway, meanwhile demonstrating the potential applications in crop biofortification. Here, we chose a tomato variety, ‘Micro-Tom’, including the WT and a mutant, *pyp1-1(H7L)*, to overexpress pepper *CCS*. Our study showed that *CCS* overexpression did not significantly affect plant growth, development, or yield (Fig. S3).

In tomato fruit, lycopene is the predominant carotenoid because the expression levels of *LCYB* are relatively limited, constraining lycopene conversion to *β*-carotene (Pecker *et al*., 1996; Ronen *et al*., 2000; Pandurangaiah *et al*., 2016). CCS has been suggested to have evolved from LCYB based on analyses of amino acid sequences (Fig. S4, (Mialoundama *et al*., 2010), sharing the same conserved motif, FLEET, and lycopene cyclase domain with LCYB (Fig. S5). Supporting this, CCS proteins were phylogenetically related to moss LCYB-like 1 and 2 proteins and other angiosperm LCYBs in the rooted phylogenetic tree (Fig. S4). The bottleneck of lycopene cyclization in tomato fruit was alleviated by CCS, attributing to cyclization of more lycopene into *β*-carotene (Fig. 4c, e). Here, the remarkable enhancement of *β*-carotene in tomato fruit is comparable or superior to previous studies overexpressing *LCYB* gene in tomato (Rosati *et al*., 2000; D’Ambrosio *et al*., 2004; Giorio *et al*., 2008; Apel & Bock, 2009; Zhu *et al*., 2020; Mi *et al*., 2022). Different from previous studies using LCYB, CCS also functioned in the conversion of antheraxanthin and violaxanthin into capsanthin and capsorubin, respectively, directing the carotenoid flux moving downstream further (Fig. 4). Besides, unlike previous studies showing transgenic plants engineered with *LCYB* exhibiting tolerance to abiotic stresses including salinity (Moreno *et al*., 2021; Mi *et al*., 2022), we observed no differences between *CCS*-overexpression lines and control lines under salinity stress condition (Fig. S6).

Xanthophyll esters in red tomato of WT and WTEV lines represented only a small proportion of carotenoids (Fig. 4g, D’Ambrosio et al., 2011). It differed from pepper since limited free xanthophylls were accumulated in tomato fruit (Fig. 1c, Fig. 4c–f). However, the increased flux in the carotenoid pathway by CCS allowed more xanthophylls to be synthesized downstream than WT and WTEV lines (Fig. 4d–f), which were available for further esterification (Fig. 4g). As a result, the proportion of xanthophyll esters in total carotenoids rose significantly from 0.74–0.81% in WT and WTEV lines to 6.55–9.89% in WTCCS lines (Fig. 4f). Collectively, total carotenoid, *β*-carotene, and xanthophylls (including zeaxanthin, antheraxanthin, (*all E*) violaxanthin, capsanthin, and capsorubin) in the *β*-branch of WTCCS fruit increased along with xanthophyll esters with positive correlation of 0.83, 0.92 and 0.87–0.96, respectively (Fig. 4, Fig. 5a), indicating the positive impact of xanthophyll esterification on carotenoid pools.

The xanthophyll esters in the mutant *pyp1-1(H7L)* were not completely missing as expected, differing from a previous study (Ariizumi *et al*., 2014). The total carotenoids and each composition, except for xanthophyll esters, were similar among WT, WTEV21, *pyp1-1(H7L)*, and 1-1EV11 lines (Fig. 3a–h, Fig. 4). However, the amounts of xanthophyll esters were at relatively low levels, accounting for 0.14–0.16% of total carotenoids in *pyp1-1(H7L)* and 1-1EV11 lines (Fig. 3h, Fig. 4g), much lower than WT and WTE21 lines. This result confirmed for the first time that the mutant *pyp1-1(H7L)* had a limited ability to esterify xanthophylls.

Furthermore, it indicated that tiny amounts of xanthophyll esters, did not have an obvious impact on the carotenoid pathway among WT, WTEV21, *pyp1-1(H7L)*, and 1-1EV11 lines (Fig. 3a–h, Fig. 4). However, when *CCS* was overexpressed in *pyp1-1(H7L)*, the xanthophyll esters were detectable but remained at relatively low levels of 0.86–1.36% of total carotenoids (Fig. 4g). Although the gene encoding XAT1 is mutated in *pyp1-1(H7L)* (Ariizumi *et al*., 2014), its paralog XAT2 (Solyc02g094430) likely remains active in this mutant. *XAT2* expression level in all tomato tissues is relatively low but slightly higher in fruit than in other tissues (CoNekT database, Proost & Mutwil, 2018), thus contributing to this limited xanthophyll esterification.

Importantly, total carotenoids of 1-1CCS lines were significantly lower than *pyp1-1(H7L)* and 1-1EV1, suggesting that in the absence of or low levels of xanthophyll esterification, the production of total carotenoids and the flux to the *β*-branch are limited (Fig. 4). Besides, it was reinforced by the correlation analysis that total carotenoids in 1-1CCS lines were negatively associated with *β*-carotene, most free xanthophylls in *β*-branch and xanthophyll esters, contrary to the alteration observed in WTCCS lines (Fig. 5).

We sought to answer why the fruit accumulated different xanthophyll esters between WT and *pyp1-1(H7L)* when *CCS* was overexpressed. In a recent study, the accumulation of free zeaxanthin was attributed to generating xanthophyll-derived apocarotenoids, which in turn mediated feedback inhibition (D’Ambrosio *et al*., 2023). This is supported by our observation that zeaxanthin amounts in 1-1CCS fruit were significantly higher than in WTCCS lines but not observed for other free xanthophylls of *β*-branch (Fig. 4f). The esterification in WTCCS lines possibly kept xanthophylls in ester form, which were stably stored in plastoglobules and avoided cleavage by CCDs (carotenoid cleavage dioxygenases) (DellaPenna and Pogson, 2006; Li et al., 2023; D’Ambrosio et al., 2023).

Though ‘Micro-Tom’ is a convenient research model, it is dwarf with small fruit (Shikata & Ezura, 2016). Hence, we selected five other large-fruited inbreds with different genetic backgrounds to test *CCS* overexpression as a strategy to potentially improve their carotenoid profiles. After crossing with ‘Micro-Tom’ WT or ‘WTCCS’, all derived F1 hybrids showed similar plant growth and fruit size to their corresponding maternal lines, indicating that the recessive dwarf allele from ‘Micro-Tom’ had no observable impact on growth and fruiting.

However, affected by introducing the genome of ‘Micro-Tom WT’ and ‘WTCCS,’ respectively, the color of fruit from control– and *CCS*-derived hybrids was altered toward red and orange, respectively, paralleling with the carotenoid compositional changes (Fig. 6). All *CCS*-derived lines showed significantly higher total carotenoid levels in fruit compared to their corresponding control-derived lines (Fig. 6b). Like the observation in WTCCS and 1-1CCS lines (Fig. 4b), lycopene in the fruits of all five *CCS*-derived varieties was depleted and converted into *β*-carotene under the catalysis of CCS (Fig. 6b). However, the accumulation of *β*-carotene in fruit varied from one *CCS*-derived hybrid to another, among which four showed significantly higher accumulation of *β*-carotene in fruit than its paternal line, ‘WTCCS’ (Fig. 6b).

In *CCS*-derived hybrids, ‘FL7907B’ x ‘WTCCS’ and ‘LA2377’ x ‘WTCCS’, the patterns in carotenoid composition were similar to ‘WTCCS’ (Fig.3, Fig. 6b). *CCS* overexpression in tomato possibly alleviated the limitation posed by the transcriptional regulation of genes in the carotenoid-associated pathway (Duduit *et al*., 2022), resulting in a higher metabolic flow moving through lycopene cyclization. Their parental lines, ‘FL7907B’, ‘LA2377’ and ‘Micro-Tom’ WT, had red fruit and accumulated dominantly lycopene but at different levels (Fig. 6b). The fruit of ‘FL7907B’ contained the highest amounts of lycopene among these three parental lines (Fig. 6b). The greater the abundance of lycopene in parental lines, the greater were the amounts of *β*-carotene in the fruit of *CCS*-derived hybrids (Fig. 6b). Consequently, ‘FL7907B’ x ‘WTCCS’ accumulated the highest *β*-carotene contents among WTCCS lines, ‘FL7907B’ x ‘WTCCS’ and ‘LA2377’ x ‘WTCCS’ hybrids (Fig. 6b).

Previously, when *LCYB* gene was expressed in tomato varieties with red fruit, including ‘Money Maker’ (Rosati *et al*., 2000; Dharmapuri *et al*., 2002), ‘Red Setter’ (D’Ambrosio *et al*., 2004), or ‘Ailsa Craig’ (Zhu *et al*., 2020), higher accumulation of *β*-carotene in fruit was observed. However, engineering carotenoid profiles in tomato with yellow or orange fruit is rare. Two varieties, ‘Juane Flamme’ and ‘LA2374’, with orange fruit and higher *β*-carotene accumulation when crossed to ‘WTCCS’ had even more improvement in *β*-carotene levels than other parental lines.

The fruit of ‘Juane Flamme’ x ‘WTCCS’ contained the highest *β*-carotene among all *CCS*-derived hybrids, which was near double that of ‘LA2374’ x ‘WTCCS’ (Fig. 6b). It was possibly caused by the different promoter region sequence of *LCYB* (or *B* allele) between ‘Juane Flamme’ and ‘LA2374’ reported previously, resulting in a higher *LCYB* expression in ‘Juane Flamme’ (Orchard *et al*., 2021). Hence, high expression of native *LCYB* in ‘Juane Flamme’ likely coordinated with CCS to produce more *β*-carotene than other *CCS*-derived hybrids (Fig. 6b). Another *CCS*-derived line, ‘LA4044’ x ‘WTCCS’, also showed a higher accumulation of *β*-carotene although it was slightly lower than that found in the fruit of ‘WTCCS’ (Fig. 6b). ‘LA4044’ has yellow fruit flesh, in which deletion of 691 bp in the promoter region of *PSY* gene resulted in its low expression level (Alseekh *et al*., 2013; Shin *et al*., 2019). It limited the flux to carotenoids since PSY catalyzes the first committed step in the pathway. Consequently, the fruit of ‘LA4044’ x ‘WTCCS’ accumulated considerably more *β*-carotene than its control-derived line but lower than the other four *CCS*-derived hybrids (Fig. 6b).

Like ‘Micro-Tom’ WT transformed with CCS, the synthesis of capsanthin in *CCS*-derived lines contributed to the increase in esterified xanthophylls (Fig. 7a). The high accumulation of total carotenoids and *β*-carotene was positively associated with the accumulation of xanthophyll esters (Fig. 4, Fig. 5). Reinforcing this, all five *CCS*-derived hybrids contained significantly higher amounts of total carotenoids and *β*-carotene (Fig. 6b), and synergistically accumulated more xanthophyll esters (Fig. 7b), compared to their corresponding control-derived hybrids. For instance, the accumulation of xanthophyll esters in the fruit of ‘LA4044’ x ‘WTCCS’ was lower than the other four *CCS*-derived lines, attributing to lower total carotenoids and *β*-carotene than the other four *CCS*-derived hybrids (Fig. 6, Fig. 7).

Apocarotenoid volatiles are known to be generated in ripe fruit via oxidation of carotenoid substates, among which *β*-ionone and *β*-cyclocitral are derived from *β*-carotene (Yoo *et al*., 2023). Apocarotenoid volatiles, including *β*-ionone, 6-methyl-5-hepten-2-one and geranylacetone, have a positive impact on fruit flavor (Klee & Tieman, 2018; Kaur *et al*., 2023) and significant natural variation for them are found in wild relatives of tomato (Barnett *et al*., 2023). Our results of volatile apocarotenoids suggested that higher accumulation of β-carotene in the fruit of *CCS*-derived hybrids contributed to higher levels of *β*-ionone and *β*-cyclocitral (Fig. 8a&b). Accordingly, lycopene-derived apocarotenoids, geranial, 6-methyl-5-hepten-2-one, 6-methyl-5-hepten-2-one and neral, showed decreased or increased or not changed depending upon the genetic background of the line (Fig. 8c–f). These results are consistent with the hypothesis that in ripe fruit of tomato plants *β*-carotene level is limiting for carotenoid cleavage dioxygenase (CCD) involved in the synthesis of β-ionone and β-cyclocitral (Yoo *et al*., 2023). The changes in volatile apocarotenoids by overexpression of *CCS* with no yield drag could inspire new strategies for improving tomato flavor, the trait most valued by the consumers (Tieman *et al*., 2017).

Retinol activity equivalent (RAE) is used to evaluate the availability of vitamin A in food (Institute of Medicine (US) Panel on Micronutrients, 2001). Unlike preformed vitamin A from animal-based foods, provitamin A, mainly *β*-carotene from plant-based foods, is a healthier form for long-term dietary intake since it requires further conversion after uptake and has not been reported to have upper limitations (Blaner, 2020). In addition, the nutritional value is higher in plant-based food in general based on nutrition density scores (Fig. S7). To evaluate the nutritional significance of *CCS*-engineered tomato lines, including ‘Micro-Tom’ (WTCCS and 1-1CCS lines) and *CCS*-derived hybrids, we summarized the RAE value from the fruit of these engineered tomato lines along with those from representative vegetables and fruits (Fig. 9).

As a widely consumed vegetable, commercial tomatoes in red, orange, and yellow colors contain limited provitamin A, much lower than many other vegetables and fruits (Fig. 9). *CCS*-engineered tomato reported here is a significantly better food source than currently available tomatoes, comparable to other high RAE vegetables and fruits or even better (Fig. 9). In particular, the RAE amounts in ripe fruit of WTCCS were comparable to carrots (Fig. 9). However, in 1-1CCS lines, the increases in RAE were constrained possibly because of the limited ability in xanthophyll esterification, so that the RAE values were lower than WTCCS lines (Fig. 9). Surprisingly, the fruit from *CCS*-derived hybrid showed even higher RAE values contributed from high *β*-carotene accumulation (Fig. 6b), among which four of them were higher than male RDA in a serving (Fig. 9), indicating the successful biofortification in different commercially viable tomato varieties. In addition to fruits and vegetables, the fruit of *CCS*-engineered tomato lines, including WTCCS, 1-1CCS, and *CCS*-derived hybrids, were comparable or even superior to other engineered crop plants in RAE value (Giuliano, 2017). For instance, compared to the first tomato (‘MoneyMaker’) engineered with *LCYB* gene from *Arabidopsis* containing ∼458 µg RAE per 100 g serving (Rosati et al., 2000), here, RAE values from the fruit of WTCCS lines and *CCS*-derived hybrids were significantly improved (Fig. 9). Golden rice contains up to 258 µg RAE per 100 g serving (Paine *et al*., 2005), which requires 271 and 349 g serving to meet female and male RDA, respectively. In this study, the consumption of 80–106 and 103–136 g of WTCCS fruit, 182–202 and 235–260 g of 1-1CCS fruit, 37–102 and 48–131 g of *CCS*-derived hybrid fruit meets female and male RDA, respectively (Fig. 9). A demographic survey documented the lack of good vitamin A consumption in 12 East African countries (Wolde & Tessema, 2023). *CCS*-engineered tomato with high provitamin A is a healthy and safe food source for combating VAD in developing nations. Besides, *CCS* overexpression in different tomato varieties could provide adequate options for catering to consumers’ demands.

In addition to provitamin A, capsanthin and capsorubin are reported to be potentially beneficial for human health (Maoka *et al*., 2001; Molnár *et al*., 2005, 2012; Fernández-García *et al*., 2016; Jo *et al*., 2017; Joo *et al*., 2021; Kennedy *et al*., 2021; Shanmugham & Subban, 2022). This study is the first report on accumulation of capsanthin and capsorubin in a fleshy fruit through metabolic engineering. Previously, ketocarotenoids were reported to accumulate in flower (*Viola cornuta* L.) and rice endosperm (Ha *et al*., 2019; Jeong *et al*., 2021). Pepper *CCS*, combined with different expression cassettes harboring carotenogenesis gene(s), was introduced into rice to generate grain accumulating up to 0.4 µg/g capsanthin and capsorubin in the endosperm (Ha *et al*., 2019). By only overexpressing the pepper *CCS* we reached comparable results. Ketocarotenoids in WTCCS and 1-1CCS fruit accumulated at higher levels (Fig. 4f) than in rice grain reported previously, while they were at similar levels in the fruits of all five F1 *CCS*-derived hybrids (Fig. 7a). Capsanthin has been reported to protect mice against hepatic steatosis and steatohepatitis in nonalcoholic fatty liver disease (Joo *et al*., 2021). A high fat diet with 0.5 µg g^−1^ capsanthin, similar to the contents in CCS-engineered fruit, generated a protective effect. (Joo *et al*., 2021).

In summary, constitutive expression of pepper *CCS* in ‘Micro-Tom’, *pyp1-1(H7L)* mutant, and selected tomato varieties with different genetic backgrounds altered the metabolic flux in the carotenoid pathway (Fig. 10). The carotenoid flux moved toward *β*-branch under the function of CCS, producing remarkably higher provitamin A, *β*-carotene (Fig. 10). Furthermore, considerable amounts of capsanthin and capsorubin accumulated in *CCS*-engineered tomato (Fig. 10). Along with the production of specialized ketocarotenoids, other xanthophylls of the *β*-branch were accumulated more in *CCS*-engineered tomato fruit, contributing to xanthophyll esterification (Fig. 10). Our results on the mutant, *pyp1-1(H7L)*, revealed that xanthophyll esters had key regulatory impacts on the carotenoid accumulation in tomato fruit, facilitating the whole carotenoid flux (Fig. 10). This study demonstrated that CCS could function under different genetic backgrounds, contributing to the accumulation of total carotenoids, *β*-carotene, ketocarotenoids and *β*-carotene-derived apocarotenoids in tomato fruit, consequently enhancing its color, flavor and nutritional value.

**Figure 10.**
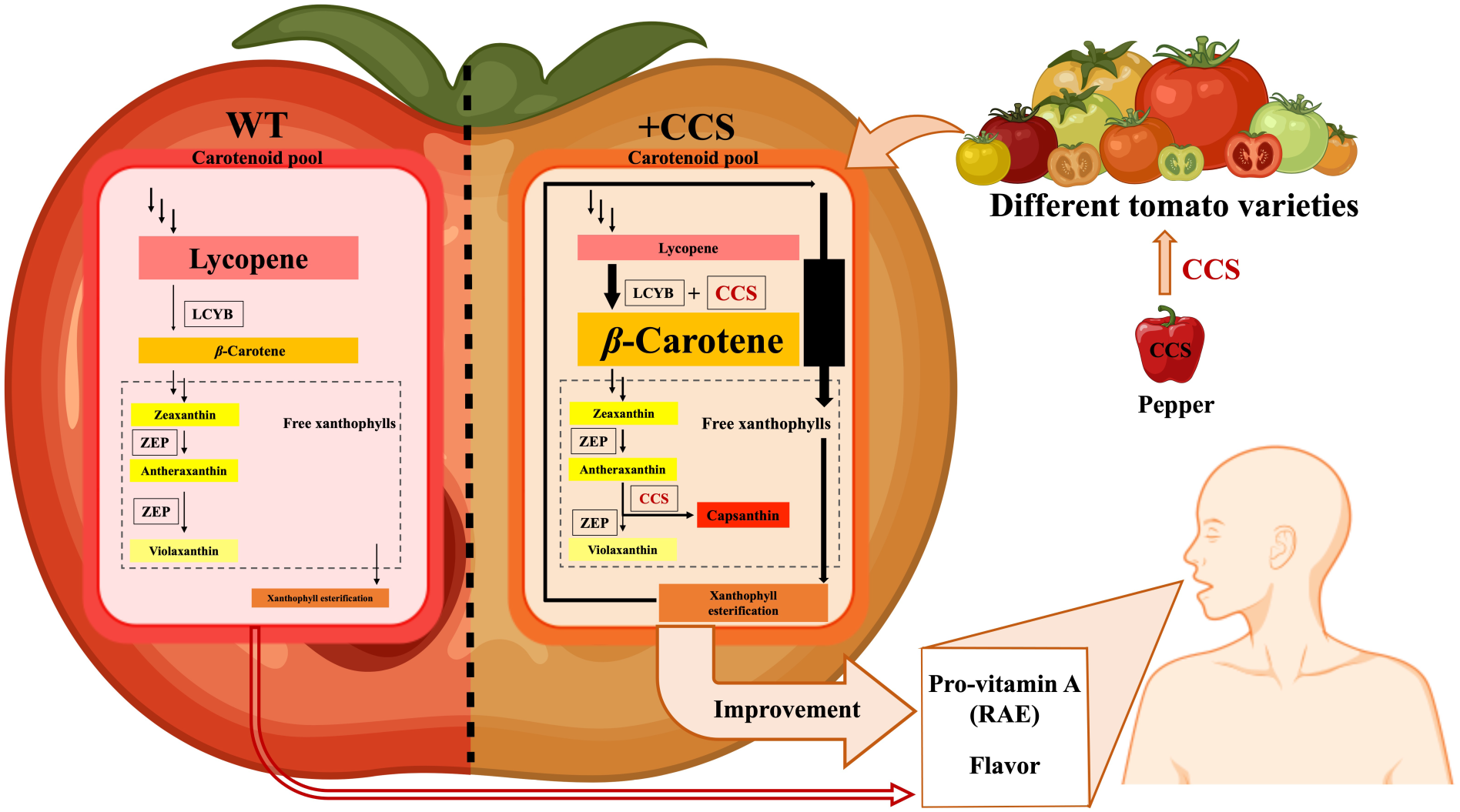
Schematic representation of phenotypic and metabolic alteration in *CCS*-transformed tomato fruit. The carotenoid pathway in WT tomato fruit is shown on the left. The altered carotenoid pathway by overexpression of pepper *CCS* in tomato fruit is shown on the right. The size of the arrow and each carotenoid represent the relative flux size and metabolite abundance in the pathway. Both pathways are shown in the presence of the xanthophyll esterification process. The accumulation of xanthophyll esters, highlighted in orange box, was positively correlated with the amounts of total carotenoids, *β*-carotene and other carotenoids, indicating its ability to facilitate carotenoid biosynthesis when *CCS* was overexpressed. Overexpression of pepper *CCS* in different tomato varieties (represented here with different color tomatoes), contributed to metabolic alterations of the carotenoid pathway and remarkably enhanced biofortification of provitamin A (*β*-carotene) for human nutrition. LCYB, lycopene *β*-cyclase; ZEP, zeaxanthin epoxidase; CCS, capsanthin/capsorubin synthase, highlighted in red. Design components of the figure were obtained from ‘BioRender’ and further assembled in PowerPoint.

## Acknowledgments

We gratefully acknowledge the funding support from the Horticultural Sciences Department, University of Florida as a graduate assistantship to J.F. B.R.’s research was supported by a grant from Florida Department of Agriculture and Consumer services via the USDA Specialty Crops Block Grant program. We gratefully acknowledge the kind donation of an *E. coli* strain, SixPack by Dr. Gyorgy Posfai and Dr. Zsuzsanna Gyorfy both from the Biological Research Centre of the Hungarian Academy of Sciences.

## Author Contributions

J.F. and B.R. planned the study, J.F. conducted the experimental work on pepper VIGS, carotenoid production in *E. coli* and transgenic tomato, D.T. conducted the work on fruit volatiles and J.F wrote the first draft, and all authors contributed to writing and editing the final draft.

## Data availability statement

All pertinent data are included in the submission.

## Supplementary Figures and Tables

**Figure S1.**
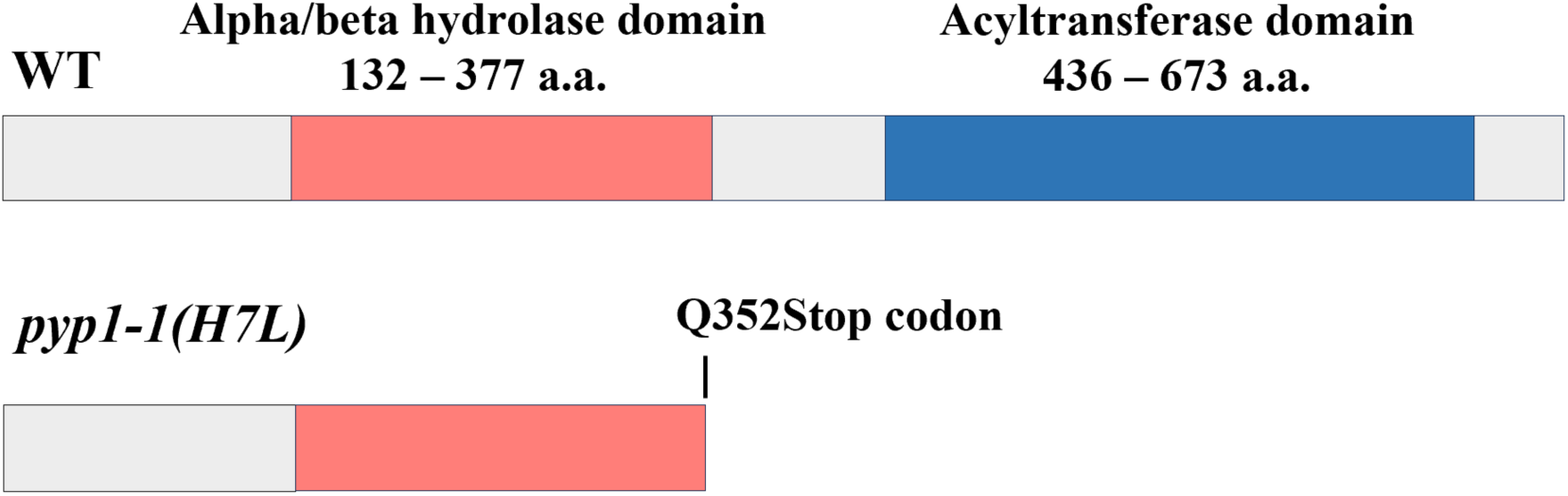
Protein structure of xanthophyll acyltransferase in WT ‘MicroTom’ (Top) and *pyp1-1(H7L)* mutant (bottom) in ‘MicroTom’ background. A non-sense mutation occurred at the 352^nd^ amino acid residue, causing a stop codon and a complete loss of the acyltransferase domain. Domain lengths are drawn not proportional to sequence length.

**Figure S2.**
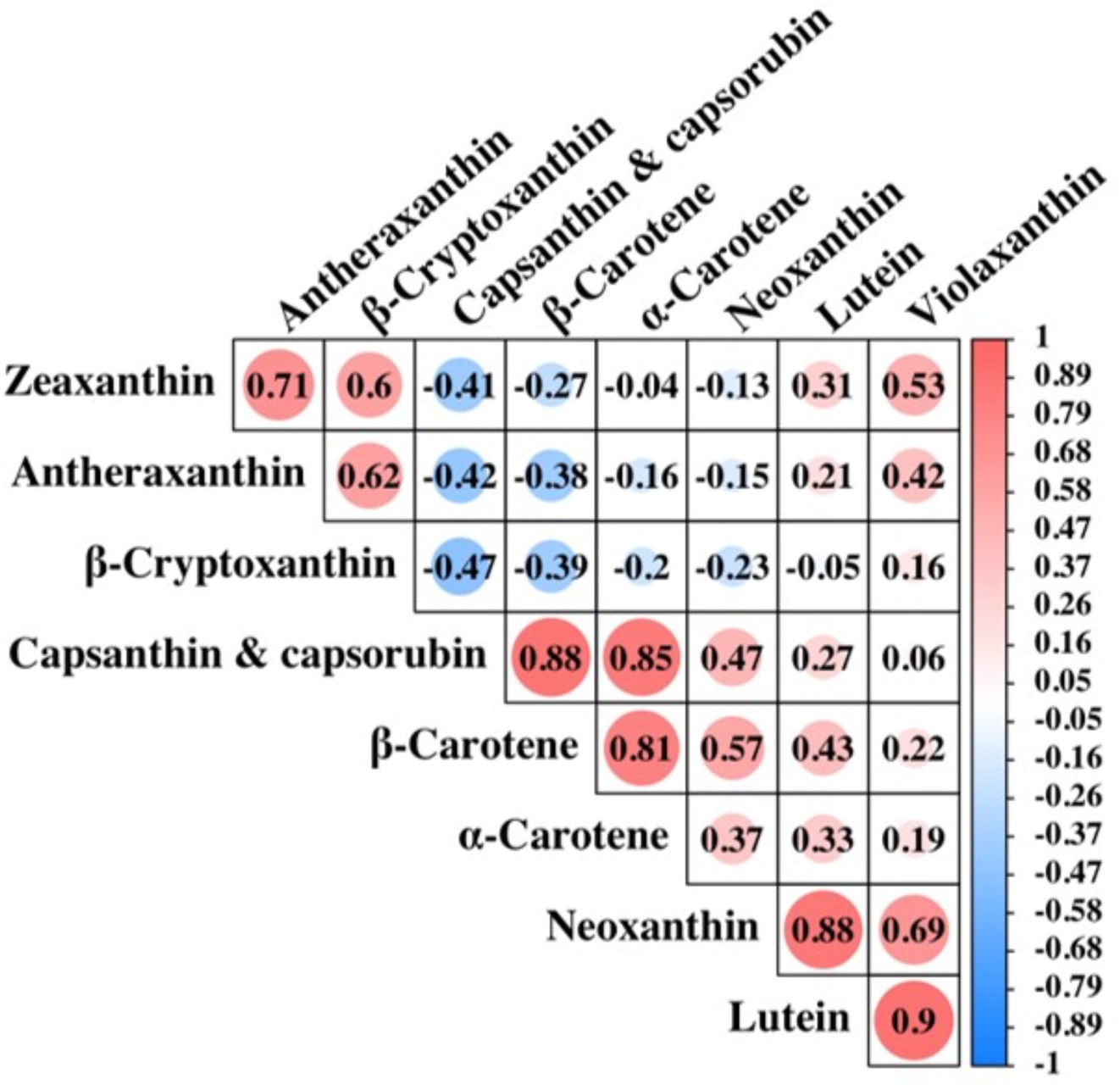
Pearson correlation matrix between each paired carotenoid content analyzed from ripe fruit of 32 pepper varieties. The data were from a previous study Wahyuni et al., (2011) and re-analyzed here. The color intensity and the size of each circle in the chart indicated a higher correlation, in which red referred to positive correlations and blue to negative correlations.

**Figure S3.**
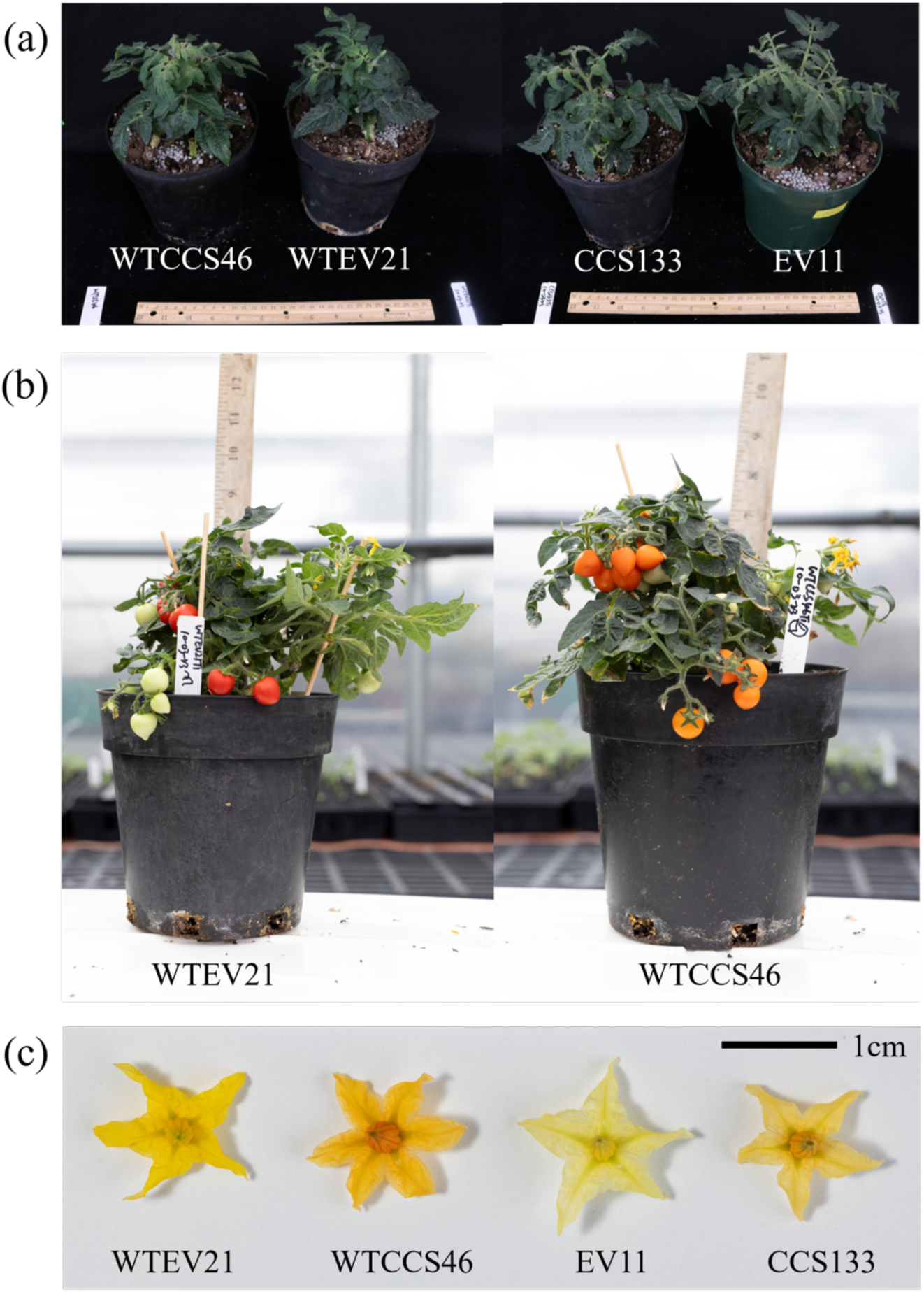
Plant growth and yield. Vegetative growth of Micro-Tom plants without or with transforming pepper *CCS* gene (a). Reproductive stage of Micro-Tom plants without or with transforming pepper *CCS* gene (b). The flower color of Micro-Tom plants without or with transforming pepper *CCS* gene (c).

**Figure S4.**
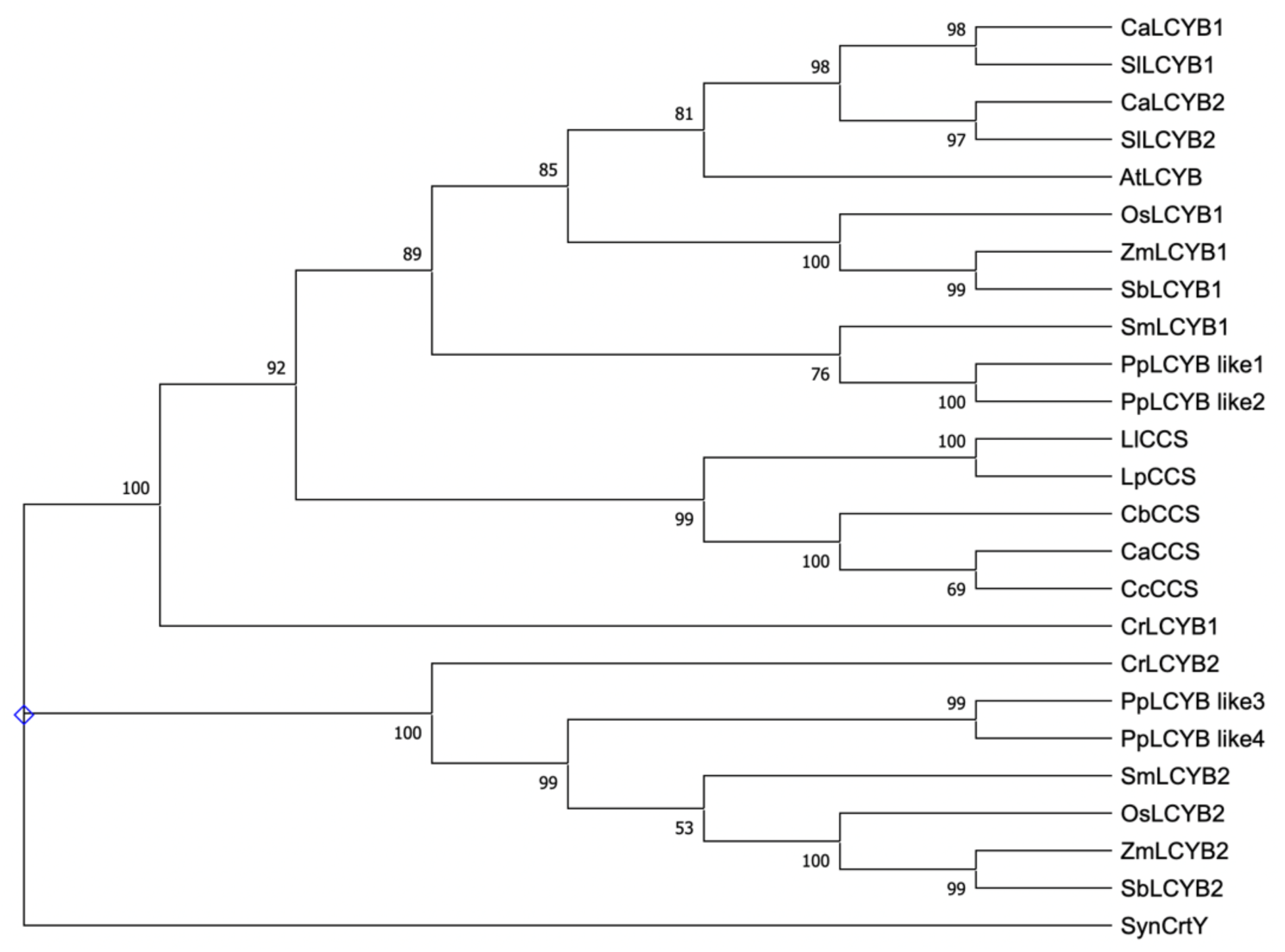
Phylogram of capsanthin/capsorubin synthase (CCS) and lycopene β-cyclase (LCYB) protein sequences. Phylogenetic analysis of CCS and LCYB protein sequences from different plant species. The rooted phylogenetic tree was generated by MEGA 11.0. CCS and LCYB protein sequences were aligned with MUSCLE program and then were constructed using the Maximum Likelihood method. The bootstrap values shown on the nodes were computed from 1000 replicates. CCS protein sequences were from *Capsicum annuum* (Ca, NP_001311998.1, Capana06g000615, CA06g22860), *Capsicum chinense* (Cc, PHU16032.1), and *Capsicum baccatum* (Cb, PHT30120.1), *Lilium lancifolium* (Ll, PHT30120.1), *Lilium pumilum* (Lp, AQZ41157.1). The LCYB protein sequence were from dicot: *Capsicum annuum* (CaLCYB1: XP_016543793; CaLYCB2: NP_001311908), *Solanum lycopersicum* (SlLCYB1: Solyc10g079480.1, XP_010312096; SlLCYB2: Solyc04g040190.1, NP_001234226), and *Arabidopsis thaliana* (AtLCYB, AT3G10230.1, NP_001078131); monocot: *Oryza sativa* (Os, OsLCYB1, XP_066163296.1; OsLCYB2, XP_015622198.1), *Zea mays* (Zm, ZmLCYB1, NP_001169155.1; ZmLCYB2, NP_001146840.1), Sorghum bicolor (SbLCYB1, XP_002453448.1; SbLCYB2, XP_002455838.1); moss: *Physcomitrella patens* (PpLCYB like1, XP_024371653.1; PpLCYB like2, XP_024373649.1; PpLCYB like3, XP_024389432.1; PpLCYB like4, XP_024380943.1); gymnosperm: *Selaginella moellendorffii* (SmLCYB1, XP_002971295.2; SmLCYB2, XP_002974344.1); Algae: *Chlamydomonas reinhardtii* (CrLCYB1, XP_042921724.1; CrLCYB2, XP_001696529.1); Cyanobacteria: *Synechococcus* sp. PCC 7335 (SynCrtY, WP_006454809.1

**Figure S5.**
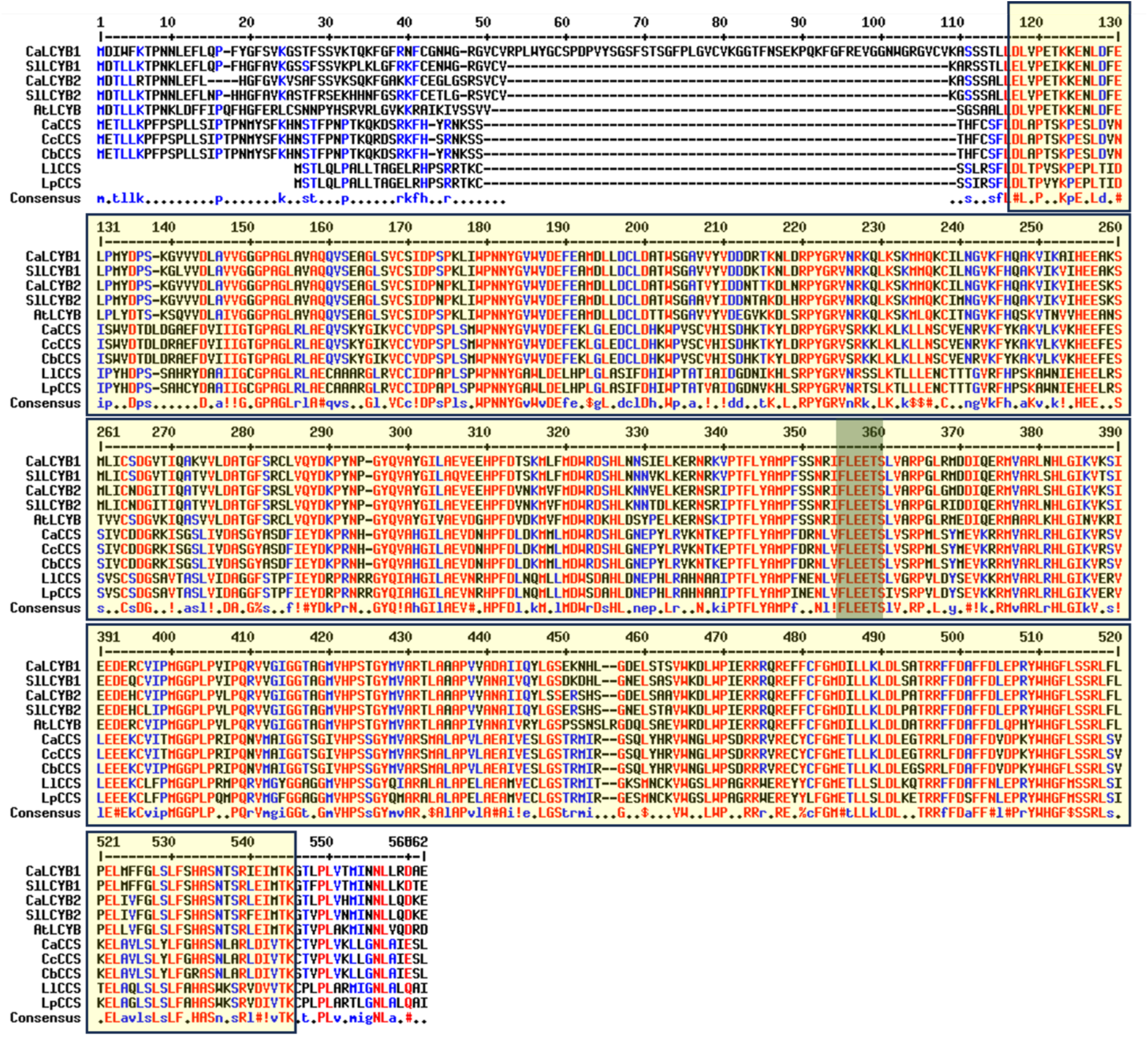
Alignment of CCS and LCYB protein sequences using MultaAlin (Corprt, 1988; http://multalin.toulouse.inra.fr/multalin/). Identical amino acids from all protein sequences were highlighted in red color. Low consensus amino acid residues from all protein sequences were highlighted in blue color. Except for CCS protein sequences mentioned in A, LCYB protein sequences were from *Capsicum annuum* (CaLCYB1: Capana10g002320, CA10g20140, XP_016543793; CaLYCB2: Capana05g000023, CA05g00080, NP_001311908), *Solanum lycopersicum* (SlLCYB1: Solyc10g079480.1, XP_010312096; SlLCYB2: Solyc04g040190.1, NP_001234226), and *Arabidopsis thaliana* (AtLCYB: AT3G10230.1, NP_001078131). The deduced amino acid sequences of CCS and LCYB contained a lycopene cyclase domain by InterPro (Paysan-Lafosse et al., 2023) and are highlighted in a yellow box. The active motif, FLEET, is highlighted in a green box.

**Figure S6.**
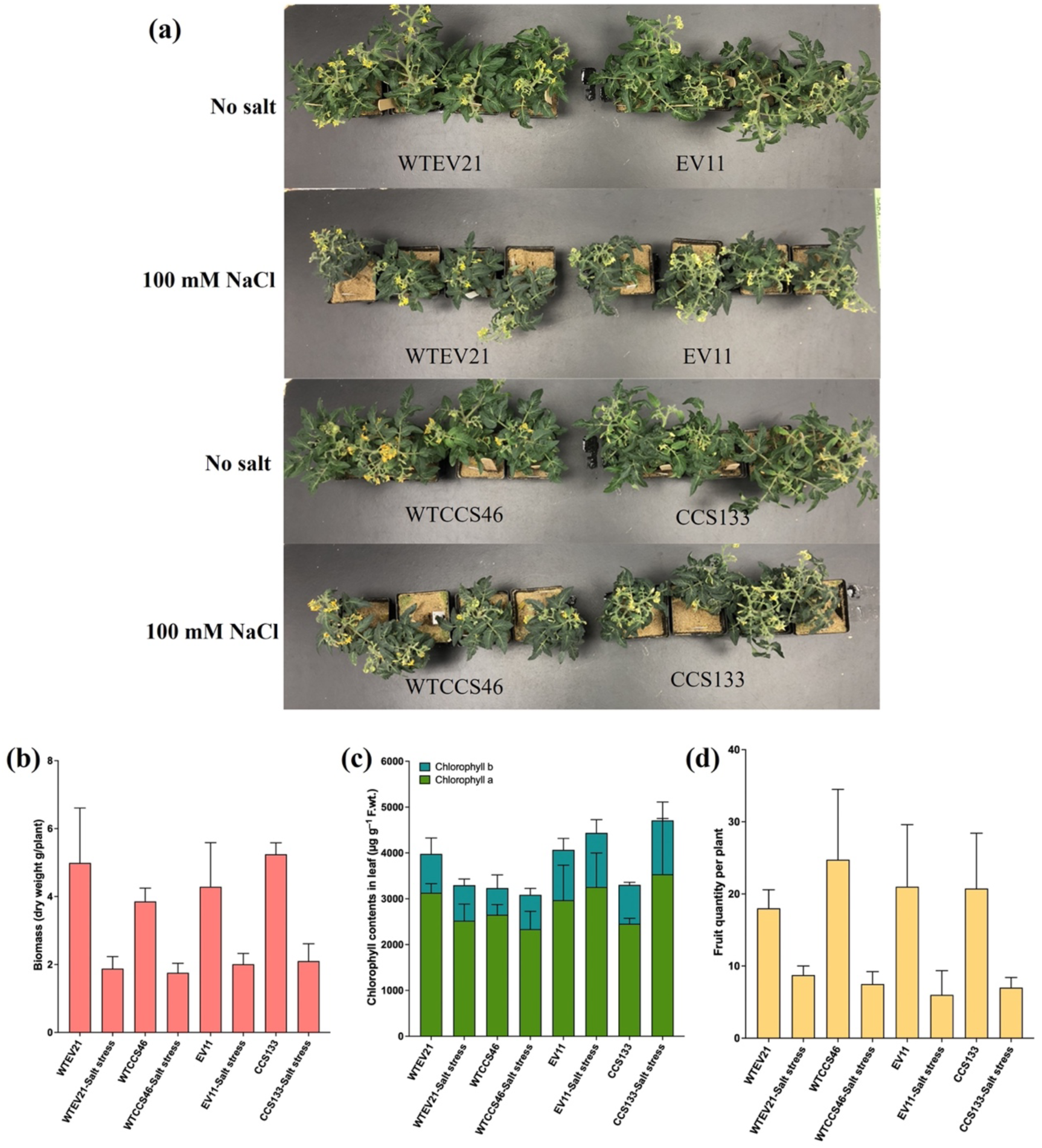
Evaluation of plant growth and salinity tolerance. Plant growth (a), plant biomass (b), chlorophyll contents (c), and fruit quantity (d) after treated with or without 100 mM NaCl. One-month-old seedlings of empty vector and *CCS*-transformed lines in Micro-Tom background or Micro-Tom *pyp1-1(H7L)* mutant background were chosen for this test. Four replicate plants at 4-5 leaf stage were transplanted in 0.5L-capacity plastic containers filled with fine vermiculite. The plants were kept under a light bench (120 µmoles.m^2^,16h light/8 h dark, 25°C/23 °C) and irrigated two pot volumes, six days per week, using half-strength Hoagland nutrient medium (pH=6.0) for control plants or with the same medium with 10 mM CaCl_2_ and NaCl (100 mM final strength, step-wise increased by 50 mM NaCl every three days and kept at 100 mM NaCl). Plants were photographed during flowering. The plants were grown for 3.5 months until the harvest of ripe fruit. Total above-ground biomass (dry weight) and fruit numbers were collected. We have not observed any developmental impacts due to CCS overexpression other than the changes in the flower and fruit color. In an experiment on container-grown plants supplied with a half-Hoagland nutrient medium, the biomass and fruit numbers were not significantly different between the vector control and CCS-overexpressing line in both wild-type and *pyp1-1(H7L)* mutant background. Also, based on biomass and fruit numbers, CCS-overexpressing lines did not have an advantage under salinity stress induced by 100 mM NaCl.

**Figure S7.**
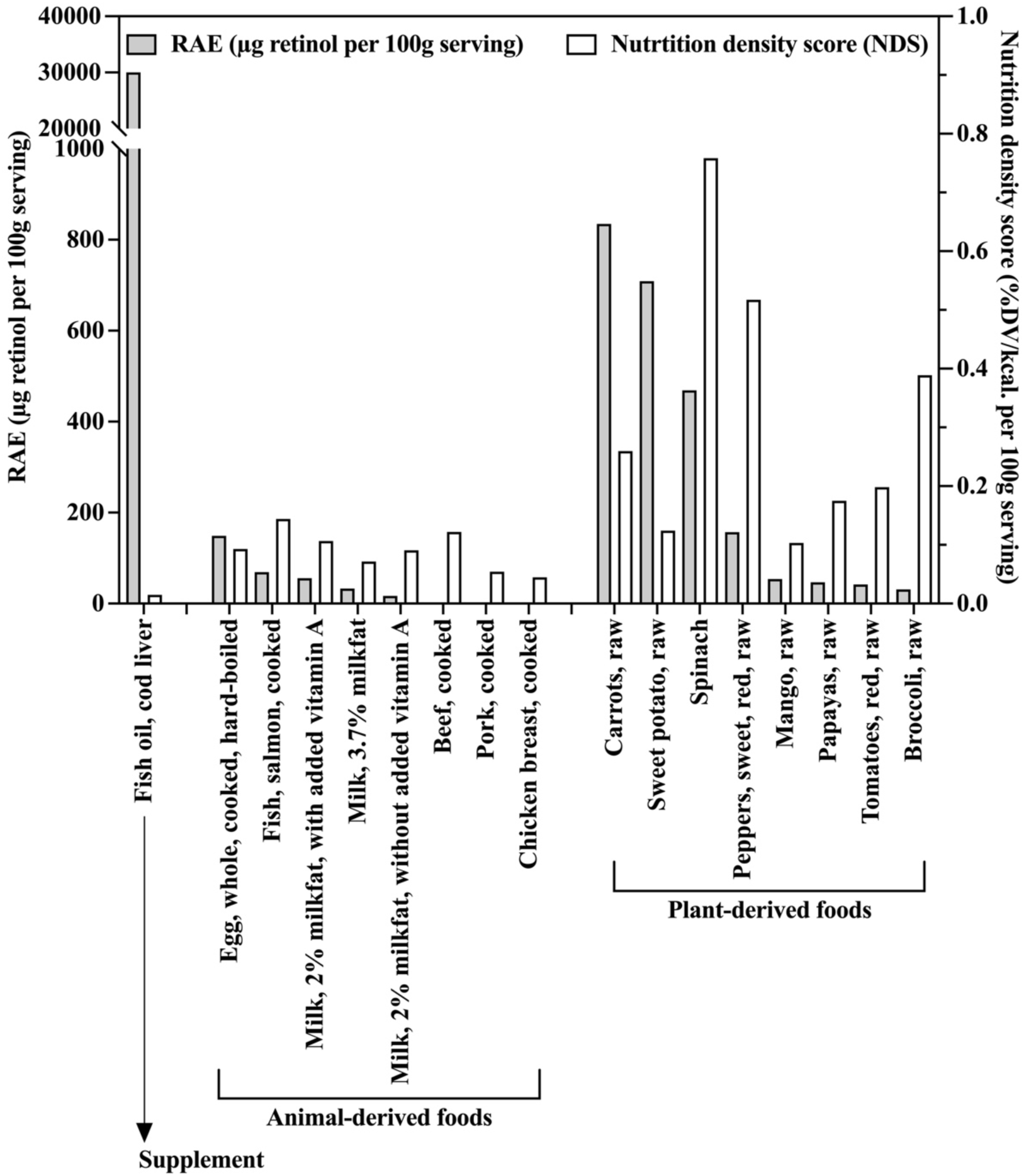
Retinol activity equivalents and nutrient density scores from multiple food sources. The data were obtained from the USDA database (standard reference legacy), the details of which were listed in a supplementary Excel file. The nutrient density scores were computed based on the method reported by Pampersaud et al., (2012).

**Figure S8.**
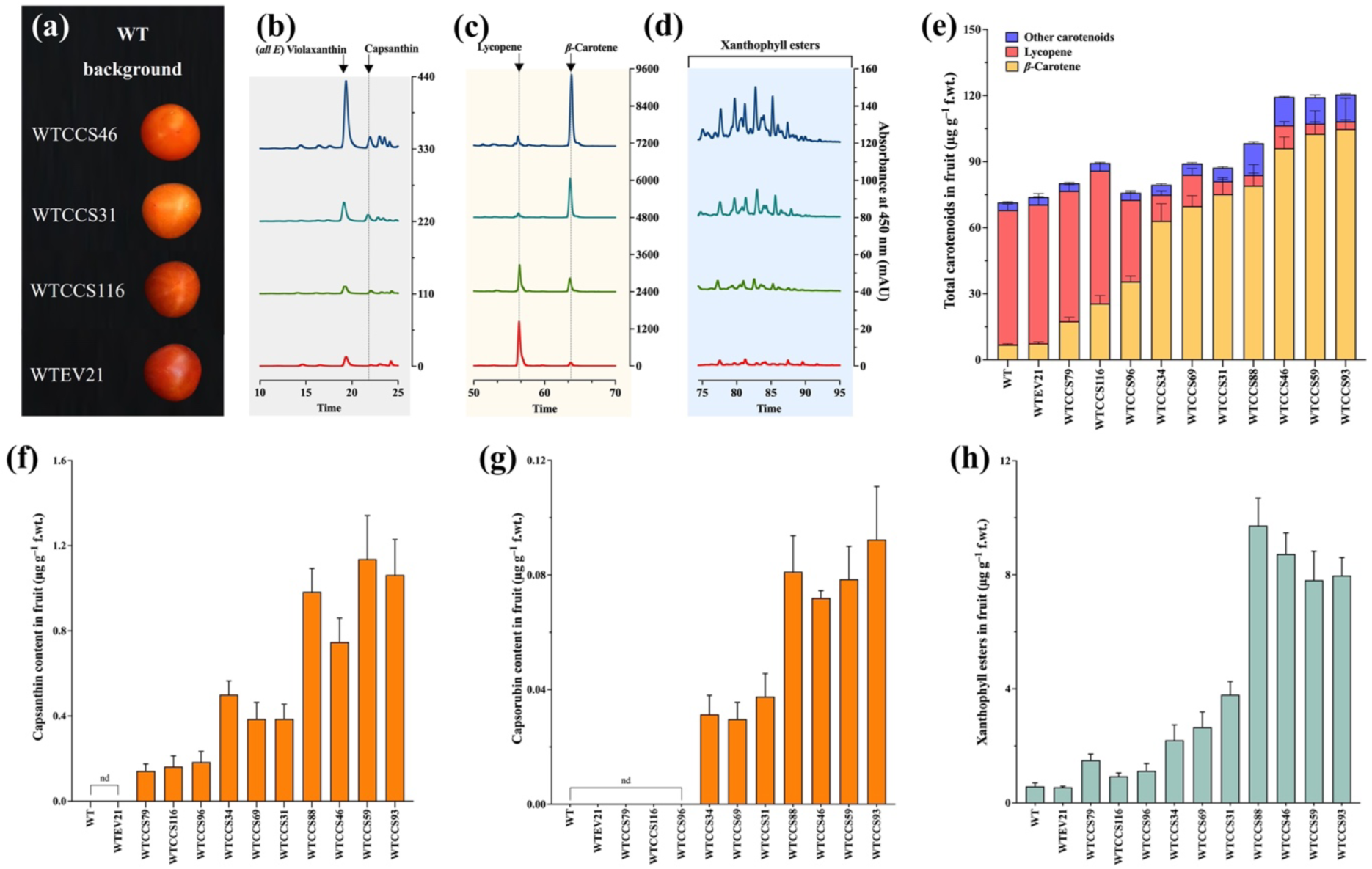
Phenotypes of the fruit from independent *CCS*-overexpression lines of tomato and their carotenoid compositions. Overexpression of pepper *CCS* was performed in ‘MicroTom’ background, including two genotypes, wildtype (WT) and *pyp1-1(H7L)* mutant for an acyltransferase defective in xanthophyll esterification. Additional independent lines potentially differing in their expression level of CCS in ‘MicroTom’ WT background were screened and analyzed. The data of WTCCS line 88, 46, 59, 93, are shown in Fig. 2 and Fig. 3, are included for comparison purposes. a. The phenotypes of tomato fruits from ‘MicroTom’ and independent *CCS*-transgenic lines in WT ‘MicroTom’ background. The fruits of independent *CCS*-transgenic lines, WTCCS116, WTCCS31 and WTCCS46, were chosen to represent low (lines 79, 116, and 96), medium (lines 34, 69, and 31), and high-(line 88, 46, 59, and 93) types classified based on fruit color and the PCA analysis on their carotenoid compositions (Fig. S6). b–d. The corresponding HPLC chromatograms to the representative fruit shown in a (WT ‘MicroTom’ background). Total carotenoids and their compositions (e), capsanthin (f), capsorubin (g), and total xanthophyll esters (h), in the fruit of independent *CCS*-transgenic lines in ‘MicroTom’ WT background. Data of I–L are presented as mean ± standard deviation of a minimum of four biological replicates based on data from HPLC analyses. One-way ANOVA was performed followed by mean separations using the Tukey test (*: P<0.05; **: P<0.01; ***: P<0.001), the details of which are shown in the supplementary Excel file.

**Figure. S9.**
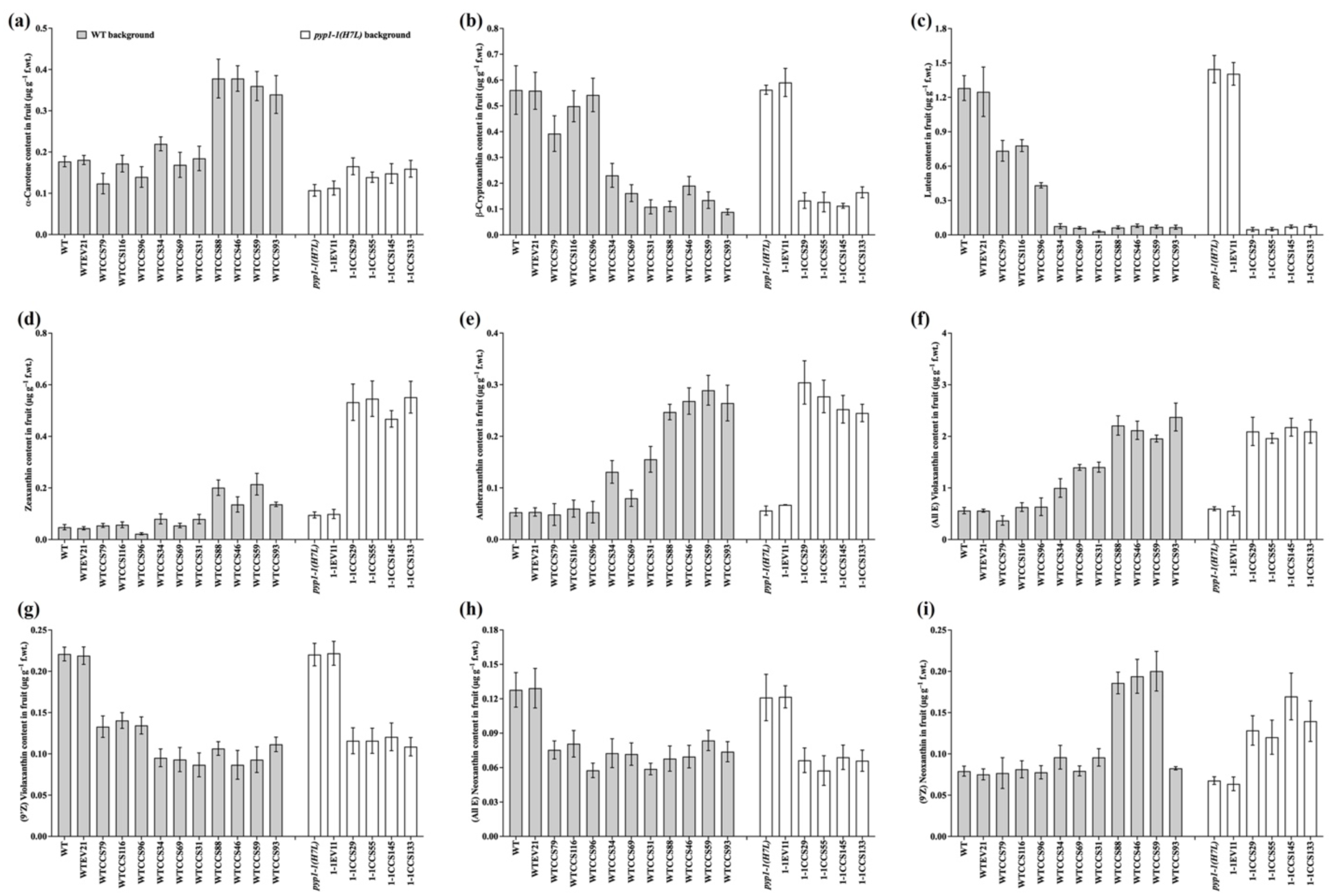
Carotenoids in the fruit of independent *CCS*-transgenic lines in ‘MicroTom’ background. Overexpression of pepper *CCS* was performed in ‘MicroTom’ background, including two genotypes, wildtype (WT) and *pyp1-1(H7L)* mutant for an acyltransferase defective in xanthophyll esterification. Multiple independent lines with different potential expression levels of CCS in ‘MicroTom’ WT background were screened and analyzed. The data of WTCCS line 88, 46, 59, 93 and 1-1CCS line 29, 55, 145, 133 shown in Fig. 2 and Fig. 3, have been included here to facilitate comparisons. Alpha-carotene (a), β-cryptoxanthin (b), lutein (c), zeaxanthin (d), antheraxanthin (e), (*all E*) violaxanthin (f), (*9’Z*) violaxanthin (g), (*all E*) neoxanthin (h), (*9’Z*) neoxanthin (i). Data are presented as mean ± standard deviation of a minimum of four biological replicates based on HPLC analyses.

**Figure S10.**
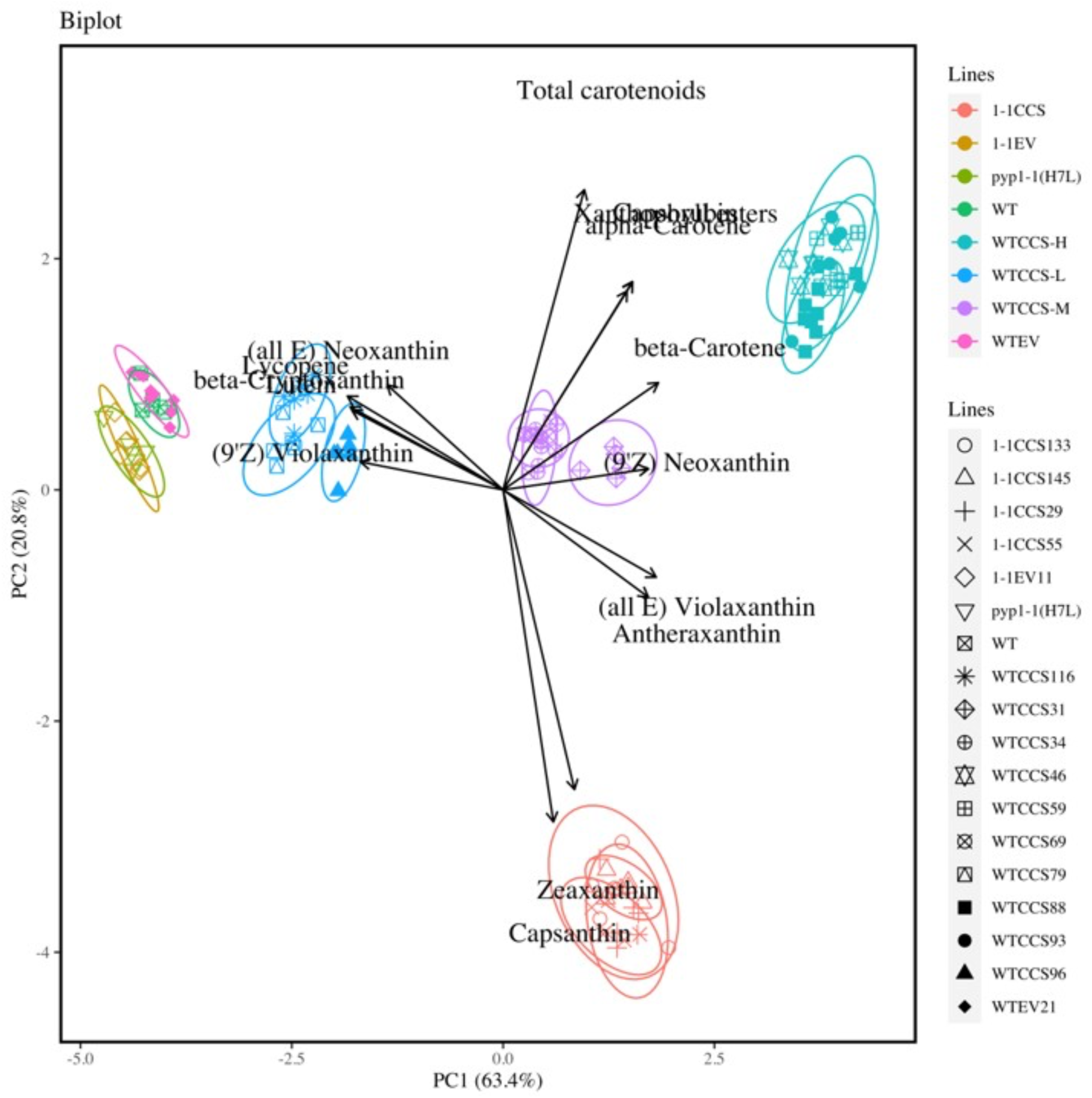
Biplot of principal component analysis (PCA) on carotenoid content in ripe tomato fruit of different *EV*-or *CCS*-transgenic lines [in both WT and *pyp1-1(H7L*) background]. PCA was performed based on the contents of carotenoids determined in fruits harvested from all lines including WT, *EV*-transformed WT, *CCS*-transformed WT, *pyp1-1(H7L)*, *EV*-transformed *pyp1-1(H7L)*, *CCS*-transformed *pyp1-1(H7L)*, which were shown in Fig. 3, Fig. S4 and Fig. S5. Biplot contained loadings, PC scores, and Proportion of variance.

**Table S1.**
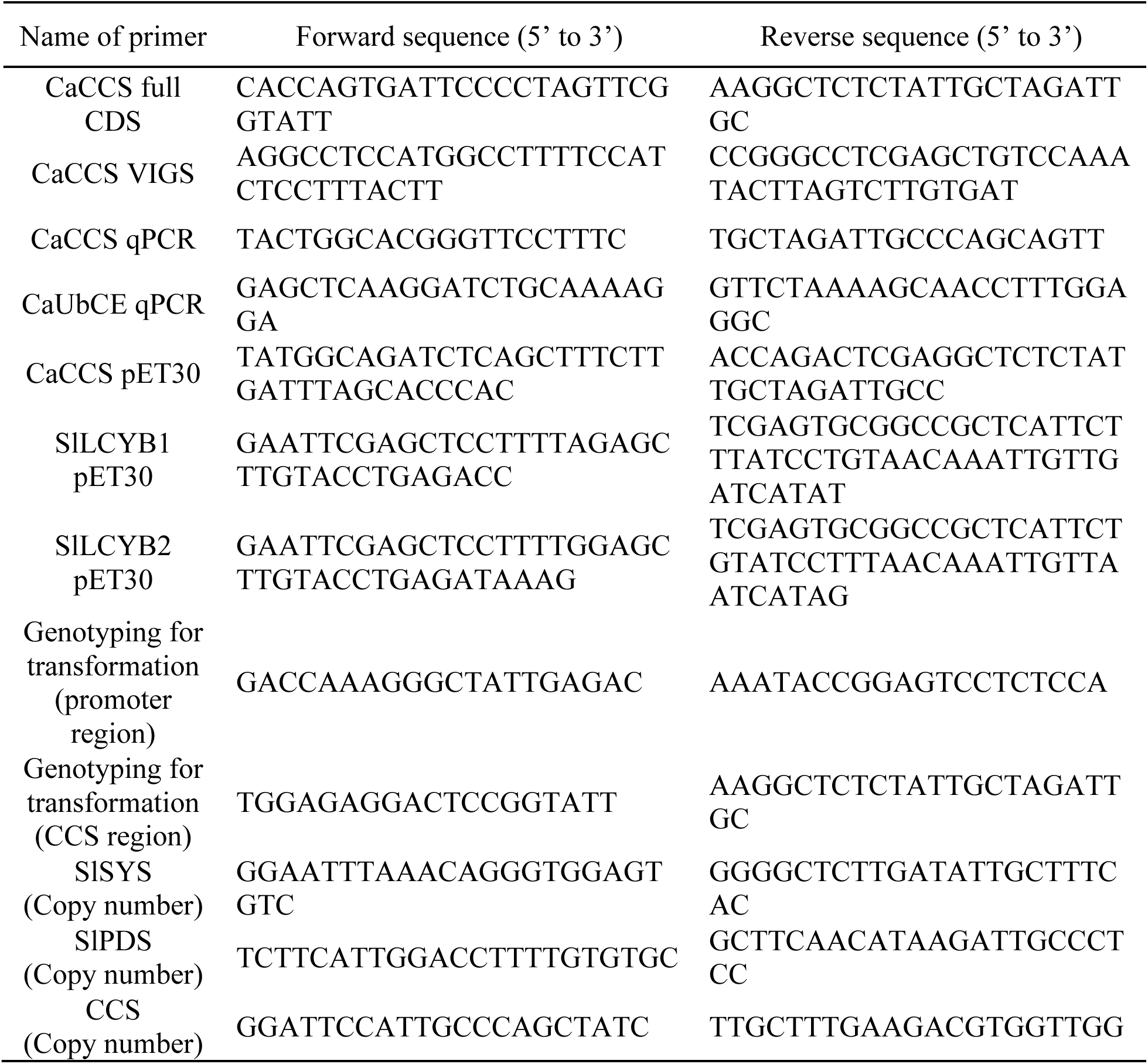
List of primers in this study.

**Table S2.** Medium for plant transformation. All media were adjusted to pH 5.7 prior to autoclaving. Plant growth regulators, antibiotic stocks, acetosyringone, and 2-mercaptoethanol, were filter-sterilized and added after autoclaving.

| Medium | Components including antibiotics, plant growth regulators and so on |
| --- | --- |
| YEP medium | YEP medium containing 50 mg L<sup>-1</sup> spectinomycin, 10 mg L<sup>-1</sup> rifampicin, and 20 mg L<sup>-1</sup> gentamycin |
| Preculture medium | MS medium supplemented with 3% (w/v) sucrose, 2 mg L<sup>-1</sup> zeatin and 0.8% (w/v) agar |
| Co-cultivation medium | MS medium supplemented with 3% (w/v) sucrose, 2 mg L<sup>-1</sup> zeatin, 100 µM acetosyringone, 10 µM 2-mercaptoethanol, and 0.8% (w/v) agar |
| Callus induction medium | MS medium supplementing with 3% (w/v) sucrose, 2 mg L<sup>-1</sup> zeatin, 100 mg L<sup>-1</sup> kanamycin, 100 mg L<sup>-1</sup> timentin, 300 mg L<sup>-1</sup> cefotaxime, and 0.8% (wt./v) agar |
| Shoot induction medium | MS medium supplemented with 3% (w/v) sucrose, 1 mg L<sup>-1</sup> zeatin, 100 mg L<sup>-1</sup> kanamycin, 100 mg L<sup>-1</sup> timentin, 300 mg L<sup>-1</sup> cefotaxime, and 0.8% (wt./v) agar |
| Rooting medium | 0.5 x MS medium supplemented with 1.5% (w/v) sucrose, 1 mg L<sup>-1</sup> IAA, 50 mg L<sup>-1</sup> kanamycin, 100 mg L<sup>-1</sup> timentin, 300 mg L<sup>-1</sup> cefotaxime, and 0.6% (w/v) agar |

**Table S3.** Chromatographic method.

| Conditions |  |
| --- | --- |
| Main column | Thermo Scientific Acclaim RSLC 120 C18 column (250 mm x 2.1 mm i.d., 2.2 μm) |
| Guard column | A similar material of main column (20 mm x 4.6 mm) |
| Temperature | 35°C |
| Injection | 1 μL |
| Flow rate | 0.1 ml/min |
| Detector | The carotenoids were detected at a wavelength of 450 nm, and the spectra were acquired and processed using the ChemStation software. |
| Mobile phase | The mobile phase consisted of eluent A (water) and eluent B (acetone) using a gradient program as follows: 75% B was increased linearly to 90% B in 20 min, subsequently holding for 15 min, then raised to 95% B in 10 min, and maintained for 20 min constantly; finally, eluent B reached 100% in 15 min and maintained for another 20 min. Initial conditions were reached in another 60 min. |

## References

1. Alseekh S, Ofner I, Pleban T, Tripodi P, Di Dato F, Cammareri M, Mohammad A, Grandillo S, Fernie AR, Zamir D. 2013. Resolution by recombination: Breaking up Solanum pennellii introgressions. Trends in Plant Science 18: 536–538.

2. Apel W, Bock R. 2009. Enhancement of carotenoid biosynthesis in transplastomic tomatoes by induced lycopene-to-provitamin a conversion. Plant Physiology 151: 59–66.

3. Ariizumi T, Kishimoto S, Kakami R, Maoka T, Hirakawa H, Suzuki Y, Ozeki Y, Shirasawa K, Bernillon S, Okabe Y, et al. 2014. Identification of the carotenoid modifying gene PALE YELLOW PETAL 1 as an essential factor in xanthophyll esterification and yellow flower pigmentation in tomato (Solanum lycopersicum). Plant Journal 79: 453–465.

4. Barnett JR, Tieman DM, Caicedo AL. 2023. Variation in ripe fruit volatiles across the tomato clade: An evolutionary framework for studying fruit scent diversity in a crop wild relative. American Journal of Botany 110: 1–14.

5. Berry HM, Rickett D V., Baxter CJ, Enfissi EMA, Fraser PD. 2019. Carotenoid biosynthesis and sequestration in red chilli pepper fruit and its impact on colour intensity traits. Journal of Experimental Botany 70: 2637–2650.

6. Blaner WS. 2020. Chapter 5 – Vitamin A and provitamin A carotenoids. In: Marriott BP, Birt DF, Stallings VA, Yates AA, eds. Present Knowledge in Nutrition (Eleventh Edition). Academic Press, 73–91.

7. Bouvier F, D’Harlingue A, Camara B. 1997. Molecular analysis of carotenoid cyclase inhibition. Archives of Biochemistry and Biophysics 346: 53–64.

8. Bouvier F, Hugueney P, D’Harlingue A, Kuntz M, Camara B. 1994. Xanthophyll biosynthesis in chromoplasts: isolation and molecular cloning of an enzyme catalyzing the conversion of 5,6-epoxycarotenoid into ketocarotenoid. The Plant Journal 6: 45–54.

9. Butelli E, Titta L, Giorgio M, Mock HP, Matros A, Peterek S, Schijlen EGWM, Hall RD, Bovy AG, Luo J, et al. 2008. Enrichment of tomato fruit with health-promoting anthocyanins by expression of select transcription factors. Nature Biotechnology 26: 1301–1308.

10. Cheng Y, Pang X, Wan H, Ahammed GJ, Yu J, Yao Z, Ruan M, Ye Q, Li Z, Wang R, et al. 2017. Identification of optimal reference genes for normalization of qPCR analysis during pepper fruit development. Frontiers in Plant Science 8: 1–14.

11. Chetty VJ, Ceballos N, Garcia D, Narváez-Vásquez J, Lopez W, Orozco-Cárdenas ML. 2013. Evaluation of four Agrobacterium tumefaciens strains for the genetic transformation of tomato (Solanum lycopersicum L.) cultivar Micro-Tom. Plant Cell Reports 32: 239–247.

12. Classic Murashige T, Skoog F. 1962. A revised medium for rapid growth and bioassays with tobacco tissue cultures. Physiol. Plant 15: 473–497.

13. Cunningham FX, Lee H, Gantt E. 2007. Carotenoid biosynthesis in the primitive red alga Cyanidioschyzon merolae. Eukaryotic Cell 6: 533–545.

14. Cunningham FX, Sun Z, Chamovitz D, Hirschberg J, Gantt E. 1994. Molecular structure and enzymatic function of lycopene cyclase from the cyanobacterium Synechococcus sp strain PCC7942. The Plant Cell 6: 1107–1121.

15. Cvetković D, Marković D. 2008. UV-effects on antioxidant activity of selected carotenoids in the presence of lecithin estimated by DPPH test. Journal of the Serbian Chemical Society 73: 1051–1061.

16. D’Ambrosio C, Giorio G, Marino I, Merendino A, Petrozza A, Salfi L, Stigliani AL, Cellini F. 2004. Virtually complete conversion of lycopene into β-carotene in fruits of tomato plants transformed with the tomato lycopene β-cyclase (tlcy-b) cDNA. Plant Science 166: 207–214.

17. D’Ambrosio C, Stigliani AL, Giorio G. 2011. Overexpression of CrtR-b2 (carotene beta hydroxylase 2) from S. lycopersicum L. differentially affects xanthophyll synthesis and accumulation in transgenic tomato plants. Transgenic Research 20: 47–60.

18. D’Ambrosio C, Stigliani AL, Rambla JL, Frusciante S, Diretto G, Enfissi EMA, Granell A, Fraser PD, Giorio G. 2023. A xanthophyll-derived apocarotenoid regulates carotenogenesis in tomato chromoplasts. Plant Science 328: 111575.

19. Delgado-Pelayo R, Hornero-Méndez D. 2012. Identification and quantitative analysis of carotenoids and their esters from sarsaparilla (Smilax aspera L.) berries. Journal of Agricultural and Food Chemistry 60: 8225–8232.

20. Dharmapuri S, Rosati C, Pallara P, Aquilani R, Bouvier F, Camara B, Giuliano G. 2002. Metabolic engineering of xanthophyll content in tomato fruits. FEBS Letters 519: 30–34.

21. Duduit JR, Kosentka PZ, Miller MA, Blanco-Ulate B, Lenucci MS, Panthee DR, Perkins-Veazie P, Liu W. 2022. Coordinated transcriptional regulation of the carotenoid biosynthesis contributes to fruit lycopene content in high-lycopene tomato genotypes. Horticulture Research 9: 1–17.

22. Fernández-García E, Carvajal-Lérida I, Pérez-Gálvez A. 2016. Carotenoids exclusively synthesized in red pepper (capsanthin and capsorubin) protect human dermal fibroblasts against UVB induced DNA damage. Photochemical and Photobiological Sciences 15: 1204–1211.

23. Fitzpatrick TB, Basset GJC, Borel P, Carrari F, DellaPenna D, Fraser PD, Hellmann H, Osorio S, Rothan C, Valpuesta V, et al. 2012. Vitamin deficiencies in humans: Can plant science help? Plant Cell 24: 395–414.

24. Furubayashi M, Kubo A, Takemura M, Otani Y, Maoka T, Terada Y, Yaoi K, Ohdan K, Misawa N, Mitani Y. 2021. Capsanthin Production in Escherichia coli by Overexpression of Capsanthin/Capsorubin Synthase from Capsicum annuum. Journal of Agricultural and Food Chemistry 69: 5076–5085.

25. Giorio G, Stigliani AL, D’Ambrosio C. 2008. Over-expression of carotene β-hydroxylase 1 (CrtR-b1) and lycopene β-cyclase (Lcy-b) in transgenic tomato fruits. Acta Horticulturae 789: 277–284.

26. Giuliano G. 2017. Provitamin A biofortification of crop plants: a gold rush with many miners. Current Opinion in Biotechnology 44: 169–180.

27. Grune T, Lietz G, Palou A, Ross AC, Stahl W, Tang G, Thurnham D, Yin S, Biesalski HK. 2010. β-Carotene Is an Important Vitamin A Source for Humans. The Journal of Nutrition 140: 2268S–2285S.

28. Guzman I, Hamby S, Romero J, Bosland PW, O’Connell MA. 2010. Variability of carotenoid biosynthesis in orange colored Capsicum spp. Plant Science 179: 49–59.

29. Ha SH, Kim JK, Jeong YS, You MK, Lim SH, Kim JK. 2019. Stepwise pathway engineering to the biosynthesis of zeaxanthin, astaxanthin and capsanthin in rice endosperm. Metabolic Engineering 52: 178–189.

30. Ha SH, Kim JB, Park JS, Lee SW, Cho KJ. 2007. A comparison of the carotenoid accumulation in Capsicum varieties that show different ripening colours: Deletion of the capsanthin-capsorubin synthase gene is not a prerequisite for the formation of a yellow pepper. Journal of Experimental Botany 58: 3135–3144.

31. Institute of Medicine (US) Panel on Micronutrients. 2001. Dietary Reference Intakes for Vitamin A, Vitamin K, Arsenic, Boron, Chromium, Copper, Iodine, Iron, Manganese, Molybdenum, Nickel, Silicon, Vanadium, and Zinc. Washington, D.C.: National Academies Press (US).

32. Jeong YS, Kim JK, Baek SA, Lee JY, Lee D, Ha SH. 2021. Reciprocal Crosses Between Astaxanthin and Capsanthin Rice Unravel Effects of Metabolic Gene Efficacy in Rice Endosperm. Journal of Plant Biology 64: 371–377.

33. Jo SJ, Kim JW, Choi HO, Kim JH, Kim HJ, Woo SH, Han BH. 2017. Capsanthin inhibits both adipogenesis in 3T3-L1 preadipocytes and weight gain in high-fat diet-induced obese mice. Biomolecules and Therapeutics 25: 329–336.

34. Joo HK, Lee YR, Lee EO, Kim S, Jin H, Kim S, Lim YP, An CG, Jeon BH. 2021. Protective Role of Dietary Capsanthin in a Mouse Model of Nonalcoholic Fatty Liver Disease. Journal of Medicinal Food 24: 635–644.

35. Kanwar P, Ghosh S, Sanyal SK, Pandey GK. 2022. Identification of Gene Copy Number in the Transgenic Plants by Quantitative Polymerase Chain Reaction (qPCR). In: Methods in Molecular Biology. Humana, New York, NY, 161–171.

36. Kaur G, Abugu M, Tieman D. 2023. The dissection of tomato flavor: biochemistry, genetics, and omics. Frontiers in Plant Science 14: 1–18.

37. Kennedy LE, Abraham A, Kulkarni G, Shettigar N, Dave T, Kulkarni M. 2021. Capsanthin, a Plant-Derived Xanthophyll: a Review of Pharmacology and Delivery Strategies. AAPS PharmSciTech 22.

38. Kim S, Ha TY, Hwang IK. 2009. Analysis, bioavailability, and potential healthy effects of capsanthin, natural red pigment from capsicum spp. Food Reviews International 25: 198–213.

39. Kim J, Park M, Jeong ES, Lee JM, Choi D. 2017. Harnessing anthocyanin-rich fruit: A visible reporter for tracing virus-induced gene silencing in pepper fruit. Plant Methods 13: 1–10.

40. Klee HJ, Tieman DM. 2018. The genetics of fruit flavour preferences. Nature Reviews Genetics 19: 347–356.

41. Kössler S, Armarego-Marriott T, Tarkowská D, Turečková V, Agrawal S, Mi J, De Souza LP, Schöttler MA, Schadach A, Fröhlich A, et al. 2021. Lycopene β-cyclase expression influences plant physiology, development, and metabolism in tobacco plants. Journal of Experimental Botany 72: 2544–2569.

42. Kraemer K, Waelti M, De Pee S, Moench-Pfanner R, Hathcock JN, Bloem MW, Semba RD. 2008. Are low tolerable upper intake levels for vitamin A undermining effective food fortification efforts? Nutrition Reviews 66: 517–525.

43. Lee PC, Schmidt-Dannert C. 2002. Metabolic engineering towards biotechnological production of carotenoids in microorganisms. Applied Microbiology and Biotechnology 60: 1–11.

44. Li R, Zeng Q, Zhang X, Jing J, Ge X, Zhao L, Yi B, Tu J, Fu T, Wen J, et al. 2023. Xanthophyll esterases in association with fibrillins control the stable storage of carotenoids in yellow flowers of rapeseed (Brassica juncea). New Phytologist.

45. Lipinszki Z, Vernyik V, Farago N, Sari T, Puskas LG, Blattner FR, Posfai G, Gyorfy Z. 2018. Enhancing the Translational Capacity of E. coli by Resolving the Codon Bias. ACS Synthetic Biology 7: 2656–2664.

46. Liu Y, Schiff M, Dinesh-Kumar SP. 2002. Virus-induced gene silencing in tomato. The Plant Journal 31: 777–786.

47. Maoka T, Fujiwara Y, Hashimoto K, Akimoto N. 2001. Capsanthone 3,6-epoxide, a new carotenoid from the fruits of the red paprika Capsicum annuum L. Journal of Agricultural and Food Chemistry 49: 3965–3968.

48. Maquilan MAD, Padilla DC, Dickson DW, Rathinasabapathi B. 2020. Improved resistance to root-knot nematode species in an advanced inbred line of specialty pepper (Capsicum annuum). HortScience 55: 1105–1110.

49. Mi J, Vallarino JG, Petřík I, Novák O, Correa SM, Chodasiewicz M, Havaux M, Rodriguez-Concepcion M, Al-Babili S, Fernie AR, et al. 2022. A manipulation of carotenoid metabolism influence biomass partitioning and fitness in tomato. Metabolic Engineering 70: 166–180.

50. Mialoundama AS, Heintz D, Jadid N, Nkeng P, Rahier A, Deli J, Camara B, Bouvier F. 2010. Characterization of plant carotenoid cyclases as members of the flavoprotein family functioning with no net redox change. Plant Physiology 153: 970–979.

51. Mínguez-Mosquera MI, Hornero-Méndez D. 1993. Separation and Quantification of the Carotenoid Pigments in Red Peppers (Capsicum annuum L.), Paprika, and Oleoresin by Reversed-Phase HPLC. Journal of Agricultural and Food Chemistry 41: 1616–1620.

52. Molnár P, Kawase M, Satoh K, Sohara Y, Tanaka T, Tani S, Sakagami H, Nakashima H, Motohashi N, Gyémánt N, et al. 2005. Biological activity of carotenoids in red paprika, Valencia orange and Golden delicious apple. Phytotherapy Research 19: 700–707.

53. Molnár J, Serly J, Pusztai R, Vincze I, Molnár P, Horváth G, Deli J, Maoka T, Zalatnai A, Tokuda H, et al. 2012. Putative supramolecular complexes formed by carotenoids and xanthophylls with ascorbic acid to reverse multidrug resistance in cancer cells. Anticancer Research 32: 507–517.

54. Moreno JC, Martinez-Jaime S, Kosmacz M, Sokolowska EM, Schulz P, Fischer A, Luzarowska U, Havaux M, Skirycz A. 2021. A Multi-OMICs Approach Sheds Light on the Higher Yield Phenotype and Enhanced Abiotic Stress Tolerance in Tobacco Lines Expressing the Carrot lycopene β-cyclase1 Gene. Frontiers in Plant Science 12: 1–17.

55. Moreno JC, Mi J, Agrawal S, Kössler S, Turečková V, Tarkowská D, Thiele W, Al-Babili S, Bock R, Schöttler MA. 2020. Expression of a carotenogenic gene allows faster biomass production by redesigning plant architecture and improving photosynthetic efficiency in tobacco. The Plant Journal: tpj.14909.

56. Neuman H, Galpaz N, Cunningham FX, Zamir D, Hirschberg J. 2014. The tomato mutation nxd1 reveals a gene necessary for neoxanthin biosynthesis and demonstrates that violaxanthin is a sufficient precursor for abscisic acid biosynthesis. Plant Journal 78: 80–93.

57. Nishino A, Yasui H, Maoka T. 2016. Reaction of Paprika Carotenoids, Capsanthin and Capsorubin, with Reactive Oxygen Species. Journal of Agricultural and Food Chemistry 64: 4786–4792.

58. Orchard CJ, Cooperstone JL, Gas-Pascual E, Andrade MC, Abud G, Schwartz SJ, Francis DM. 2021. Identification and assessment of alleles in the promoter of the Cyc-B gene that modulate levels of β-carotene in ripe tomato fruit. Plant Genome 14.

59. Paine JA, Shipton CA, Chaggar S, Howells RM, Kennedy MJ, Vernon G, Wright SY, Hinchliffe E, Adams JL, Silverstone AL, et al. 2005. Improving the nutritional value of Golden Rice through increased pro-vitamin A content. Nature Biotechnology 23: 482–487.

60. Pandurangaiah S, Ravishankar K V., Shivashankar KS, Sadashiva AT, Pillakenchappa K, Narayanan SK. 2016. Differential expression of carotenoid biosynthetic pathway genes in two contrasting tomato genotypes for lycopene content. Journal of Biosciences 41: 257–264.

61. Pecker I, Gabbay R, Cunningham FX, Hirschberg J. 1996. Cloning and characterization of the cDNA for lycopene beta-cyclase from tomato reveals decrease in its expression during fruit ripening. Plant Molecular Biology 30: 807–819.

62. Proost S, Mutwil M. 2018. CoNekT: An open-source framework for comparative genomic and transcriptomic network analyses. Nucleic Acids Research 46: W133–W140.

63. Ronen G, Carmel-Goren L, Zamir D, Hirschberg J. 2000. An alternative pathway to β-carotene formation in plant chromoplasts discovered by map-based cloning of Beta and old-gold color mutations in tomato. Proceedings of the National Academy of Sciences of the United States of America 97: 11102–11107.

64. Rosati C, Aquilani R, Dharmapuri S, Pallara P, Marusic C, Tavazza R, Bouvier F, Camara B, Giuliano G. 2000. Metabolic engineering of beta-carotene and lycopene content in tomato fruit. Plant Journal 24: 413–420.

65. Saito T, Ariizumi T, Okabe Y, Asamizu E, Hiwasa-Tanase K, Fukuda N, Mizoguchi T, Yamazaki Y, Aoki K, Ezura H. 2011. TOMATOMA: A novel tomato mutant database distributing micro-tom mutant collections. Plant and Cell Physiology 52: 283–296.

66. Shanmugham V, Subban R. 2022. Capsanthin from Capsicum annum fruits exerts anti-glaucoma, antioxidant, anti-inflammatory activity, and corneal pro-inflammatory cytokine gene expression in a benzalkonium chloride-induced rat dry eye model. Journal of Food Biochemistry 46: 1–16.

67. Shikata M, Ezura H. 2016. Micro-Tom Tomato as an Alternative Plant Model System: Mutant Collection and Efficient Transformation. In: Methods in Molecular Biology. 47–55.

68. Shin JH, Yoo HJ, Yeam I, Lee JM. 2019. Distinguishing two genetic factors that control yellow fruit color in tomato. Horticulture Environment and Biotechnology 60: 59–67.

69. Van Der Straeten D, Bhullar NK, De Steur H, Gruissem W, MacKenzie D, Pfeiffer W, Qaim M, Slamet-Loedin I, Strobbe S, Tohme J, et al. 2020. Multiplying the efficiency and impact of biofortification through metabolic engineering. Nature Communications 11: 1–10.

70. Sun HJ, Uchii S, Watanabe S, Ezura H. 2006. A highly efficient transformation protocol for Micro-Tom, a model cultivar for tomato functional genomics. Plant and Cell Physiology 47: 426–431.

71. Tian SL, Li L, Shah SNM, Gong ZH. 2015. The relationship between red fruit colour formation and key genes of capsanthin biosynthesis pathway in Capsicum annuum. Biologia Plantarum 59: 507–513.

72. Tieman DM, Zeigler M, Schmelz EA, Taylor MG, Bliss P, Kirst M, Klee HJ. 2006. Identification of loci affecting flavour volatile emissions in tomato fruits. Journal of Experimental Botany 57: 887–896.

73. Tieman D, Zhu G, Resende MFR, Lin T, Nguyen C, Bies D, Rambla JL, Beltran KSO, Taylor M, Zhang B, et al. 2017. A chemical genetic roadmap to improved tomato flavor. Science (New York, N.Y.) 355: 391–394.

74. Trajković M, Jevremovic S, Dragićević M, Simonović AD, Subotić AR, Miloševi S, Cingel A. 2021. Alteration of flower color in viola Cornuta cv. “lutea splendens” through metabolic engineering of Capsanthin/Capsorubin synthesis. Horticulturae 7.

75. Wahyuni Y, Ballester A-R, Sudarmonowati E, Bino RJ, Bovy AG. 2011. Metabolite biodiversity in pepper (Capsicum) fruits of thirty-two diverse accessions: Variation in health-related compounds and implications for breeding. Phytochemistry 72: 1358–1370.

76. Wahyuni Y, Ballester AR, Sudarmonowati E, Bino RJ, Bovy AG. 2013. Secondary metabolites of Capsicum species and their importance in the human diet. Journal of Natural Products 76: 783–793.

77. Wolde M, Tessema ZT. 2023. Determinants of good vitamin A consumption in the 12 East Africa Countries using recent Demographic and health survey. PLoS ONE 18: 1–15.

78. Xu S, Gao S, An Y. 2023. Research progress of engineering microbial cell factories for pigment production. Biotechnology Advances 65: 108150.

79. Ye X, Beyer P. 2000. Engineering the provitamin A (β-carotene) biosynthetic pathway into (carotenoid-free) rice endosperm. Science 287: 303–305.

80. Yoo HJ, Chung MY, Lee HA, Lee S Bin, Grandillo S, Giovannoni JJ, Lee JM. 2023. Natural overexpression of CAROTENOID CLEAVAGE DIOXYGENASE 4 in tomato alters carotenoid flux. Plant Physiology 192: 1289–1306.

81. Zheng X, Giuliano G, Al-Babili S. 2020. Carotenoid biofortification in crop plants: citius, altius, fortius. Biochimica et Biophysica Acta – Molecular and Cell Biology of Lipids 1865.

82. Zhu K, Zheng X, Ye J, Jiang Q, Chen H, Mei X, Wurtzel ET, Deng X. 2020. Building the synthetic biology toolbox with enzyme variants to expand opportunities for biofortification of provitamin a and other health-promoting carotenoids. Journal of Agricultural and Food Chemistry 68: 12048–12057.

